# The histone lysine demethylase KDM5C fine-tunes gene expression to regulate dendritic cell heterogeneity and function

**DOI:** 10.1101/2023.05.28.542441

**Authors:** Hannah Guak, Matthew Weiland, Alexandra Vander Ark, Lukai Zhai, Kin Lau, Batsirai Mabvakure, Mario Corrado, Paula Davidson, Shelby Compton, Lisa DeCamp, Catherine Scullion, Russell G. Jones, Sara M. Nowinski, Connie M. Krawczyk

## Abstract

The functional and phenotypic heterogeneity of dendritic cells (DCs) plays a crucial role in facilitating the development of diverse immune responses that are essential for providing host protection. We found that KDM5C, a histone <u>lysine</u> <u>d</u>e<u>m</u>ethylase of the KDM5 family regulates several aspects of conventional DC (cDC) and plasmacytoid DC (pDC) population heterogeneity and function. Using mice conditionally deficient in KDM5C in DCs, we found that loss of KDM5C results in an increase in Ly6C^−^ pDCs compared to Ly6C^+^ pDCs. We found that Ly6C^−^ pDCs, compared to Ly6C^+^ pDCs, have increased expression of cell cycle genes, decreased expression of activation markers and limited ability to produce type I interferon (IFN). Both KDM5C-deficient Ly6C^−^ and Ly6C^+^ pDCs have increased expression of activation markers, however, are dysfunctional and have limited ability to produce type I IFN. For conventional cDCs, KDM5C deficiency resulted in increased proportions of cDC2Bs (CLEC12A^+^, ESAM^−^) and cDC1s, which was partly dependent on type I IFN and pDCs. Using ATAC-seq, RNA-seq, and CUT&RUN for histone marks, we found that KDM5C regulates epigenetic programming of cDC1. In the absence of KDM5C, we found an increased expression of inflammatory markers, consistent with our previous results in bone marrow-derived DCs. However, we also found a decrease in mitochondrial metabolism genes and altered expression of cDC lineage-specific genes. In response to *Listeria* infection, KDM5C-conditionally deficient mice mounted reduced CD8^+^ T cell responses, indicating that KDM5C expression in DCs is necessary for their function. Thus, KDM5C is a key regulator of DC heterogeneity by modulating the balance of DC subsets and serves as a critical driver of the epigenetic programming and functional properties of DCs.

## Introduction

Dendritic cells (DCs) are innate immune cells that play key roles in shaping innate and adaptive immunity. Like many immune cell types, DCs are a heterogeneous population comprising subsets that are classified based on ontogeny, transcriptional signatures, and functional properties. Their functional and phenotypic heterogeneity enables them to orchestrate customized immune responses that afford host protection against a diverse range of threats. Conventional or classical DCs (cDCs) are myeloid-derived and are functionally divided into two major subsets: cDC1 (CD8^+^XCR1^+^) and cDC2 (CD4^+^CD172a^+^CD11b^+^). cDC2 can be further divided into cDC2A (ESAM^+^CLEC12A^−^) and cDC2B (ESAM^−^CLEC12A^+^), which are defined by the expression of the transcription factors T-BET and RORγt, respectively (1). cDC subsets are generally thought to arise from a common pool of pre-DCs that are committed to preferentially differentiate into cDC1 or cDC2 (2,3). cDCs are exceptionally efficient at presenting antigen to T cells, with cDC1s being specialized to cross-present antigens and stimulate CD8^+^ T cell responses, whereas cDC2s promote CD4^+^ T cell responses.

Another class of DC known as plasmacytoid dendritic cells (pDCs) can participate in T cell priming in some contexts, but they are primarily known for their capacity to produce large amounts of type I interferons (IFNs). Although less phenotypic heterogeneity has been described for pDCs, there is increased evidence of heterogeneity in the pDC population (4–6). In addition, pDC-like cells that possess characteristics of both pDCs and cDCs have been described in mice and humans (7–9). These cells have a transcriptional profile associated with pDCs but are poor producers of type I IFN and have an increased capacity for antigen presentation. Recent studies have used single-cell RNA-seq to better delineate DC ontogeny and have shown that pDCs are predominantly lymphoid in origin, although the precise contribution from the myeloid lineage is still debated (4,10,11). In fact, one recent study suggests that pDCs and cDC1s have a common precursor that is distinct from cDC2 precursors (11), and another identifies a pDC-like population that can give rise to cDC2Bs (12). Thus, the origins and differentiation trajectory of pDCs are still actively being defined, along with the mechanisms that guide DC fate.

Transcription factors such as Interferon Regulatory Factor (IRF) 8 and IRF4 are transcription factors that promote lineage specificity and function of pDCs and cDCs. IRF8 is required for the development of cDC1s as well as for the maintenance of cDC1 identity once differentiated (13). Although IRF8 is not required for pDC differentiation, it is essential for pDC function, including for type I IFN production (13). In contrast, IRF4 is not required for cDC1 or pDC differentiation and function but supports those of a subset of cDC2s (14,15). The amount of IRF8 or IRF4 is key for cDC identity, as high amounts of IRF8 is required for cDC1 identity, and high abundance of IRF4 can induce a similar transcriptional program including the majority of cDC1-specific genes (16).

Several transcriptional regulators in addition to IRF8 and IRF4 support epigenetic and transcriptional programming controlling DC specification (1,17–19). Deletion of these factors leads to either reduced production of a specific DC subset and/or a DC subset with abnormal identity (15,20–22). For example, deletion of the cDC1-specific factor BATF3 leads to the development of a cDC1-like subset that also displays cDC2-like properties (20). BATF3 maintains IRF8 autoactivation in cDC1-committed progenitors to support optimal cDC1 development (21). As well, production of cDC2B from pDC-like cells is dependent on KLF4 (12). Together these studies demonstrate the key role that transcription factors have in DC specification and function, but the role of chromatin modifiers such as KDM5C in these processes is poorly understood.

We previously found that the histone <u>lysine d</u>e<u>m</u>ethylase KDM5C (SMCX, JARID1C) restrains bone marrow-derived DC (BMDC) activation *in vitro*, which was the first evidence that KDM5C is an important regulator of immune cell function (23). KDM5C removes permissive methyl groups from H3K4, thereby acting as a transcriptional repressor (24). In some instances, however, KDM5C has been associated with transcriptional activation (25,26). *Kdm5c* is an X-linked gene that escapes X-inactivation, so females express more KDM5C than do males(27). While a function for KDM5C in immune cells has not been described, KDM5C has been shown to regulate immune genes in non-immune cell types (28–31).

To better understand the function of KDM5C in DCs *in vivo,* we generated mice deficient in KDM5C in DCs (*Kdm5c^ΔItgax^* and *Kdm5c^ΔZbtb46^*). We found that in the absence of KDM5C, pDC and cDC heterogeneity is altered. *Kdm5c^ΔItgax^*mice have an increase in Ly6C^−^ pDCs, which are poor producers of type I IFN. Both KDM5C-deficient Ly6C^−^ and Ly6C^+^ pDCs are also impaired in producing type I IFN. Additionally, *Kdm5c^ΔItgax^* mice display a specific increased proportion of cDC1 compared to cDC2, accompanied by an imbalance in cDC2 subsets, with a marked increase in cDC2Bs and decrease in cDC2A. Mechanistically, we found that KDM5C regulates epigenetic and transcriptional programming, leading to enhanced expression of inflammatory genes despite reduced DC function. Mitochondrial metabolism was altered in cDC1, and the expression of cDC1 lineage-specific genes including *Irf8* was reduced in KDM5C-deficient cDC1. *Kdm5c^ΔItgax^* mice had reduced CD8^+^ T cell responses to *Listeria* infection, demonstrating the requirement of KDM5C for cDC1 function. Together, our data show that the histone lysine demethylase KDM5C uniquely functions in DCs to regulate DC specificity and function.

## Results

### KDM5C regulates pDC heterogeneity

Previously, we have shown that KDM5C cooperates with PCGF6 to restrain BMDC steady-state activation (23). To investigate the function of KDM5C in DCs *in vivo*, we generated mice with KDM5C deleted in pDC and cDCs (*Itgax-Cre-Kdm5c^fl/fl^; Kdm5c^ΔItgax^*), and examined the abundance and proportions of DC populations in the spleen (**Figs. 1, S1**). Within the pDC population (Lineage(B220^+^CD3^+^Ly6C^+^CD19^+^NK1.1^+^) SiglecH^+^CD11c^int^CD11b^−^PDCA1^+^), we discovered that a significant proportion of splenic pDCs in the *Kdm5c^ΔItgax^*mice did not express Ly6C. This Ly6C^−^ population was increased > 4-fold compared to the controls (**Fig 1A**). The proportion of splenic Ly6C^+^ pDCs was lower in *Kdm5c^ΔItgax^* due to the increase in Ly6C^−^ pDCs, although the cell count was unchanged (**Fig 1A**).

**Figure 1 -.**
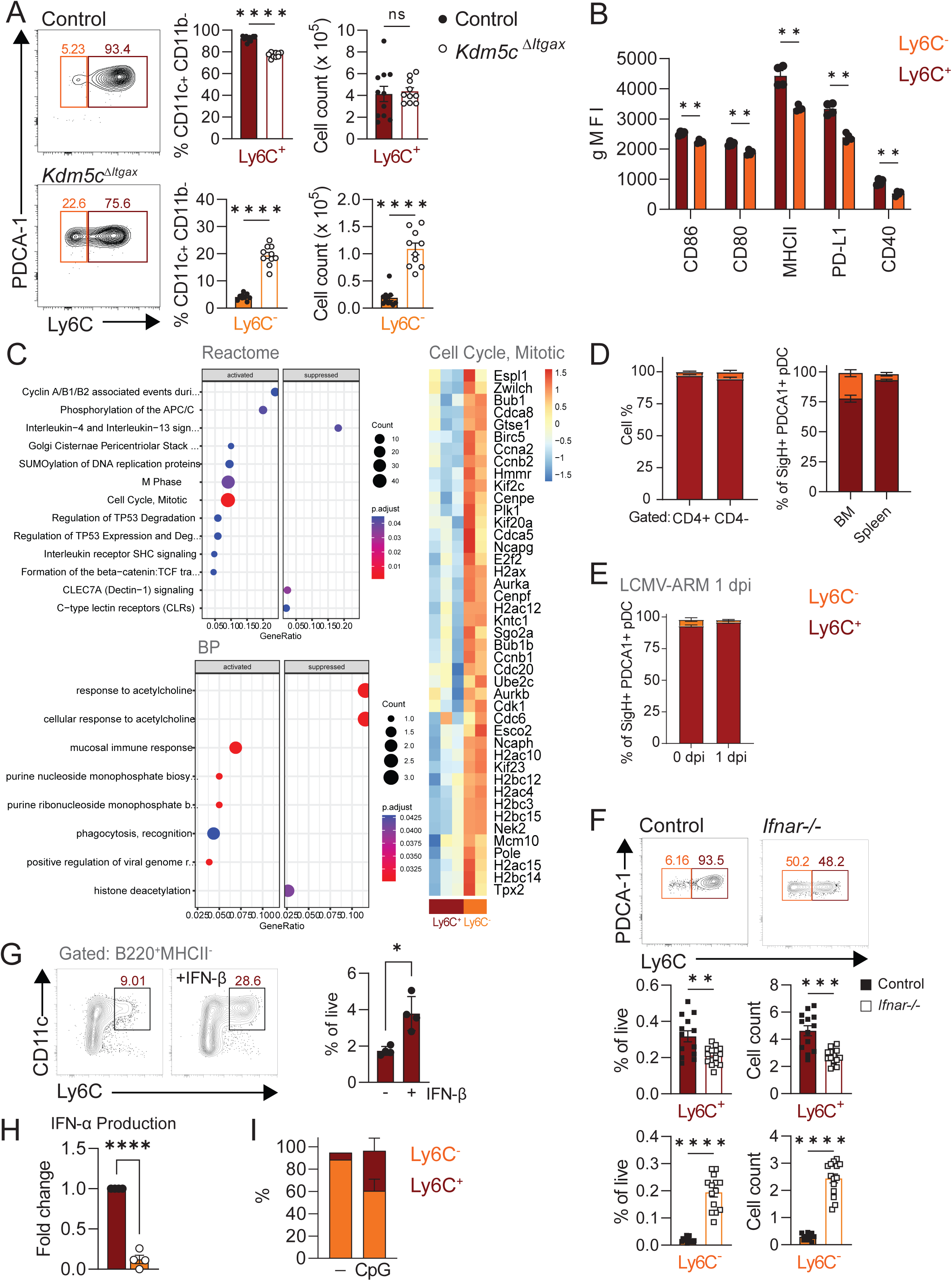
KDM5C regulates pDC heterogeneity. (**A**) The proportions and cell counts of splenic Ly6C^+^ pDCs and Ly6C^−^ pDCs from female control and *Kdm5c^ΔItgax^* mice were determined by flow cytometry. Numbers above gates in representative FACS plots (left) indicate frequencies of Ly6C^+^ pDCs or Ly6C^−^ pDCs as a percentage of the parent population (Lin(B220)^+^SiglecH^+^CD11c^int^CD11b^−^). (**B**) Geometric mean fluorescence intensity (MFI) of activation markers CD80, MHCII, PD-L1, CD86 and CD40 expressed by Ly6C^+^ (red bars) and Ly6C^−^ pDCs (orange bars). (**C**) Dot plots of RNA-seq analysis comparing Ly6C^+^ and Ly6C^−^ pDCs with Reactome Pathway Analysis and GSEA of Biological Processes (left); heatmap of cell cycle genes (right). (**D**) Proportions of CD4^+^ and CD4^−^ cells as a percentage of Ly6C^+^ pDCs (dark red bars) or Ly6C^−^ pDCs (orange bars) (left) and percentage of Ly6C^+^ or Ly6C^−^ pDCs of Lin(B220)^+^SiglecH^+^CD11c^int^CD11b^−^ cells in bone marrow (BM) and spleen (right). (**E**) Proportions of splenic pDCs at steady state and 1 day post infection with LCMV Armstrong (LCMV-ARM). (**F**) Frequencies and cell counts of splenic Ly6C^+^ pDCs and Ly6C^−^ pDCs from WT and *Ifnar1*^−/−^ mice. Numbers above gates in representative FACS plots indicate percentages of Ly6C^+^ pDCs or Ly6C^−^ pDCs of live cells. (**G**) Proportion of Ly6C^+^CD11c^+^B220^+^MHCII^−^ pDCs as a percentage of live cells following differentiation of bone marrow with FLT3L in the presence of IFN-β (50 U/mL). (**H**) IFN-α production by sorted splenic Ly6C^+^ and Ly6C^−^ pDCs stimulated with TLR9 ligand CpG ODN 1585 for 18 hours, measured by ELISA and expressed as fold change relative to stimulated control Ly6C^+^ pDCs. (**I**) Percentages of Ly6C^+^ and Ly6C^−^ pDCs of total cells following CpG stimulation of sorted Ly6C^−^ pDCs. Each symbol from (**A, B, H**) represents an individual mouse. Data are (**A**) pooled from 2 experiments (mean and s.e.m. of 10 to 11 mice per group), (**B**) one experiment representative of three experiments (mean and s.e.m. of 4 mice per group), (**F**) one experiment representative of three experiments (mean and s.d. of four replicates per group), (**G, H**) one experiment (mean and s.e.m. of 4 biological replicates per group), (**I**) pooled from three experiments (mean and s.e.m. of 13 to 14 mice per group). Statistical significance was determined by unpaired *t*-test. * p < 0.05, ** p < 0.01, *** p < 0.001, **** p < 0.0001

To further describe Ly6C^−^ pDCs, we examined activation marker expression and found that Ly6C^−^ pDCs have significantly lower expression of the surface markers typically associated with DC activation, including CD86, CD80, MHCII, PD-L1, and CD40, compared to Ly6C^+^ pDCs (**Fig 1B**). We examined transcriptional differences between Ly6C^+^ and Ly6C^−^ pDCs using RNA-seq and found that Ly6C^−^ pDCs have increased enrichment of genes associated with the cell cycle, whereas Ly6C^+^ pDCs have increased enrichment of C-type lectins and IL-4 and IL-13 signaling (**Fig 1C**). While pDCs are terminally differentiated and are known to have low proliferation potential (32), these data suggest that the Ly6C^−^ pDCs still have proliferative capacity and are less mature. CD4^−^ pDCs have been shown to be a less mature subset of pDC (32); we therefore examined whether Ly6C expression correlates with CD4 expression on pDCs. Ly6C^−^ and CD4-expressing cells do not co-segregate; however, Ly6C^+^ pDCs have a greater proportion of CD4^+^ cells compared to Ly6C^−^ pDCs (**Fig 1D, left**). Further, bone marrow, where pDC development takes place, contains a larger frequency of Ly6C^−^ pDCs compared to those found in the spleen (**Fig 1D, right**). These data suggest Ly6C^−^ pDCs are less mature than Ly6C^+^ pDCs.

To determine if pDC populations change following immune activation, we examined pDC populations in uninfected controls and mice infected with lymphocytic choriomeningitis virus (LCMV) for 20 hours. We found that very few Ly6C^−^ pDCs were present in the spleen. (**Fig 1E**). Because type I IFN feedback is known to be important for pDC function (33), we tested whether type I IFN is required for the steady state populations of Ly6C^−^ and Ly6C^+^ pDC by examining mice deficient for IFNAR1 (*Ifnar1^−/−^*), the receptor for IFN-α and IFN-β. *Ifnar1^−/−^* mice had less splenic Ly6C^+^ pDCs and more Ly6C^−^ pDCs compared to the wild-type (WT) controls (**Fig 1F**). In line with these results, the generation of Ly6C^+^ pDCs was enhanced by the addition of IFN-β during *in vitro* differentiation of bone marrow precursors to DCs with FLT3L (**Fig 1G**). To test the functionality of Ly6C^−^ pDC, we measured their ability to produce type I IFN following TLR9 stimulation. We found that Ly6C^−^ pDC produced less IFN-α compared to Ly6C^+^ pDCs (**Fig 1H**). As well, about half of the Ly6C^−^ pDC became Ly6C^+^ following TLR9 stimulation (**Fig 1I**). Together, these findings suggest that type I IFN affects the balance of Ly6C^−^/^+^ pDCs and that Ly6C^−^ pDCs may be a less mature subpopulation of pDCs.

To determine how KDM5C deficiency impacts pDC phenotype, we examined activation markers by flow cytometry. KDM5C-deficient animals had increased expression of the activation markers CD80, MHC-II, and CD40 in both Ly6C^+^ and Ly6C^−^ pDCs, and increased PD-L1 expression by Ly6C^+^ pDCs, compared to controls (**Fig 2A**). We generated bone marrow chimeras to test whether the proportions of Ly6C^−^ and Ly6C^+^ pDC are due to intrinsic factors or can be rescued by a WT environment and/or WT pDCs. We reconstituted the bone marrow of CD45.1 mice (WT) with either an equal mix of CD45.1^+^ control bone marrow cells and CD45.2^+^ *Kdm5c^ΔItgax^* cells (KO), or of CD45.2^+^ *Kdm5c^ΔItgax^* cells alone. In a WT environment with and without WT pDC, *Kdm5c^ΔItgax^* bone marrow produced similar proportions of Ly6C^−^ and Ly6C^+^ pDCs compared to *Kdm5c^ΔItgax^*mice (**Fig 2B**). Similarly, activation markers expressed on KO Ly6C^+/-^ pDCs that developed in a WT environment remained at levels of *Kdm5c^ΔItgax^* alone (**Fig 2B**). Therefore, we conclude that Ly6C^−^ pDC population size is intrinsically regulated by KDM5C.

**Figure 2 -.**
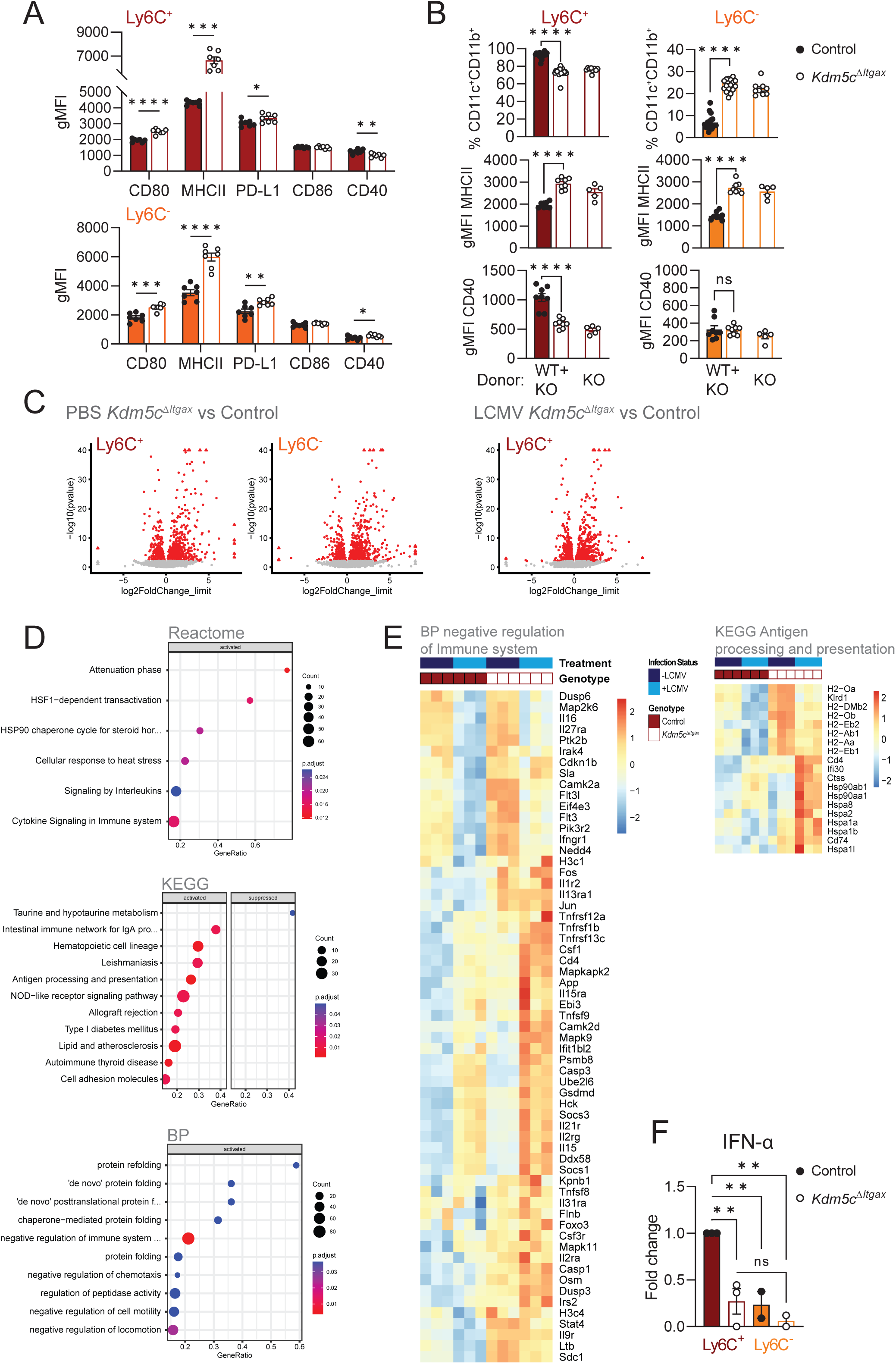
KDM5C alters transcriptional profiles and impairs type I IFN production by pDCs. (**A**) Geometric MFI of activation markers CD80, MHCII, PD-L1, CD86, and CD40 of Ly6C^+^ and Ly6C^−^ pDCs from control or *Kdm5c^ΔItgax^*mice. (**B**) Bone marrow cells that were either a 1:1 mixture of CD45.1^+^ WT and CD45.2^+^ *Kdm5c^ΔItgax^* cells or CD45.2^+^ *Kdm5c^ΔItgax^* cells alone were transferred into lethally irradiated CD45.1^+^ WT mice. Proportions of splenic cDC subsets derived from CD45.1^+^ WT and CD45.2^+^ *Kdm5c^ΔItgax^*cells and geometric MFI of MHCII and CD40 were determined 7 weeks post-injection. (**C**) Volcano plots of RNA-seq of control and *Kdm5c^ΔItgax^*Ly6C^−^ and Ly6C^+^ pDCs with or without LCMV infection (20 hours). Red data points indicate differential expression. (**D**) Significantly different Pathway and GSEA analysis of control and Kdm5c*^ΔItgax^*Ly6C^+^ pDCs with LCMV infection. (**E**) Heat maps of genes from select pathways from (**D**); BP pathway negative regulation of immune system (left) and KEGG pathway antigen processing and presentation (right). (**F**) IFN-ɑ production by sorted splenic Ly6C^+^ and Ly6C^−^ pDCs from control and *Kdm5c^ΔItgax^* mice that were stimulated with TLR9 ligand CpG ODN 1585 for 18 hours, measured by ELISA and expressed as fold change relative to stimulated control Ly6C^+^ pDCs. Data are of (**A**) one experiment representative of > three experiments (mean and s.e.m. of 7 mice per group), (**B**) pooled from two experiments (cell proportions; mean and s.e.m. of 18 mice from mixed donor group and 9 mice from *Kdm5c^ΔItgax^* donor alone) or one experiment representative of two experiments (activation markers; mean and s.e.m. of 8 mice for mixed donor group and 5 mice for *Kdm5c^ΔItgax^*donor alone), (**F**) pooled averages from two to three experiments (mean and s.e.m. of 2 to 3 per group, with each data point representing an individual experiment). Statistical significance was determined by (**A**) unpaired *t-*test, or (**B, F**) one-way ANOVA. * p < 0.05, ** p < 0.01, *** p < 0.001, **** p < 0.0001

We examined how KDM5C deficiency affects transcriptional programming in pDCs by comparing the transcriptomes of control and KDM5C-deficient Ly6C^−^ and Ly6C^+^ pDCs from mice injected with PBS or LCMV (**Fig 2C**). In response to LCMV (20 hours), KDM5C-deficient pDCs had increased expression of genes involved in antigen presentation, interferon signaling (IFI genes), and cytokine signaling, suggesting increased immune activation (**Fig 2D, E and S2**). We also found enrichment of genes involved in negative regulation of the immune system (**Fig 2E**). Because we found increased gene expression of genes that both positively and negatively regulate immune responses, we tested the function of KDM5C-deficient pDCs. In response to TLR9 stimulation, KDM5C-deficient Ly6C^+^ pDCs produced significantly less IFN-ɑ compared to controls (**Fig 2F**). Thus, in the absence of KDM5C, pDCs have an activated transcriptional profile and cell surface phenotype but fail to produce type I IFN upon stimulation.

### KDM5C regulates cDC heterogeneity

Our initial DC population profiling experiments included cDCs (Lineage^−^CD64^−^MHCII^+^CD11c^+^ CD26^+^) at homeostasis and we discovered that loss of KDM5C also impacts cDC heterogeneity. The proportion and number of cDC1s (XCR1^+^) and cDC2Bs (CD172a^+^ESAM^−^CLEC12A^+^) and merocytic DCs (XCR1^−^CD172a^−^) were higher in *Kdm5c^ΔItgax^*mice compared to control mice (**Fig 3A**), whereas cDC2As (ESAM^+^CLEC12A^−^) were lower (1,5,34,35). The proportional changes were largely due to a decrease in cDC2A numbers **(Supplementary Fig 3A**).

**Figure 3 -.**
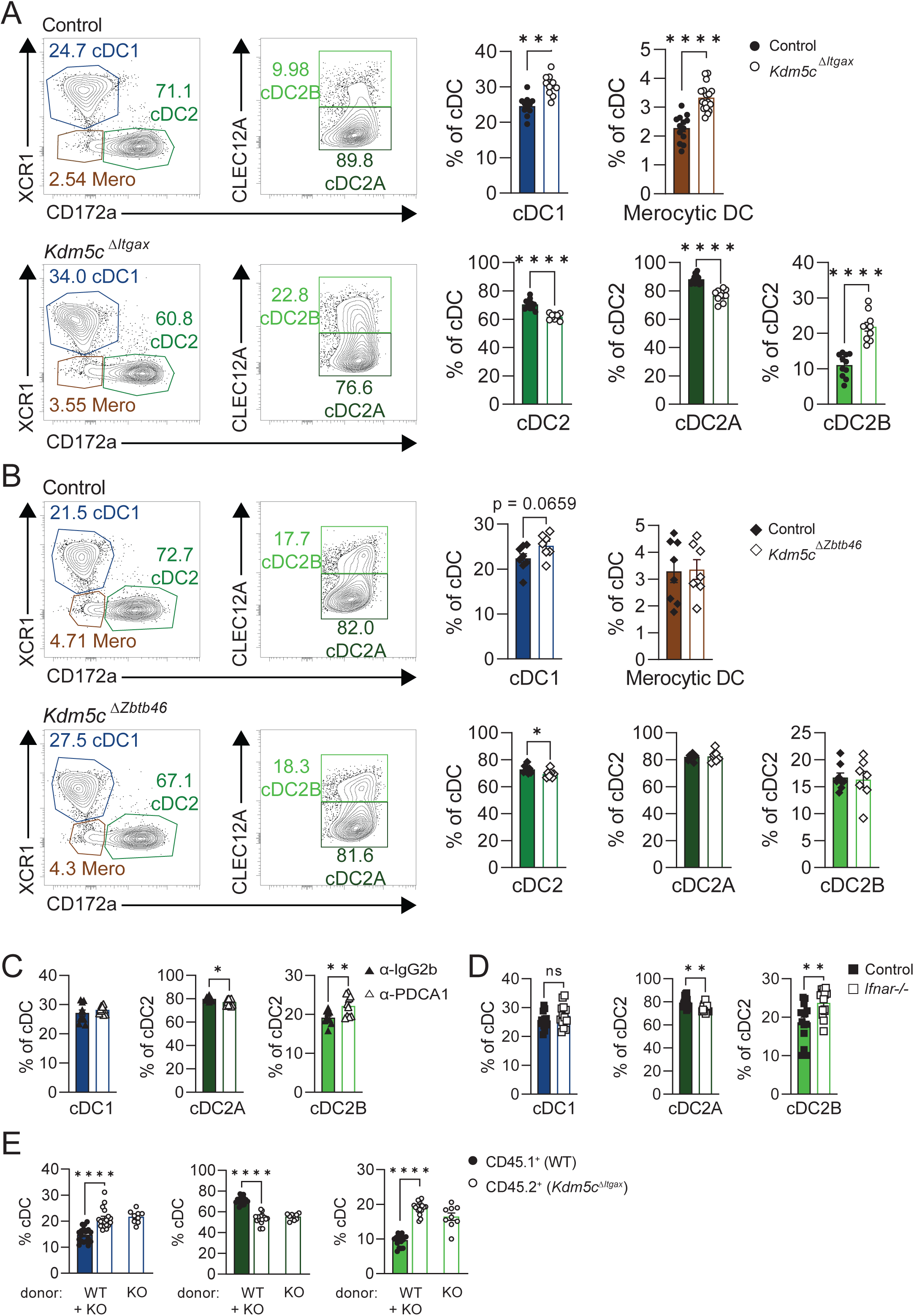
KDM5C regulates cDC heterogeneity. In female control and *Kdm5c^ΔItgax^* mice, the cell counts and frequencies of (**A**) splenic XCR1^+^ cDC1s, XCR1^−^CD172a^−^ merocytic DCs (top), CLEC12A^−^ cDC2A, and CLEC12A^+^ cDC2B (bottom) were determined by flow cytometry. Numbers next to gates in representative FACS plots (left) indicate percentages of cDC1, cDC2, or merocytic DCs as a percentage of total cDCs (Lin^−^MHCII^+^CD11c^+^CD26^+^) or cDC2A or cDC2B as a percentage of CD172a^+^ cDC2s. (**B**) As in (**A**), cDC subset proportions were determined by flow cytometry in *Kdm5c^ΔZbtb46^*. Proportions of cDC1, cDC2A, and cDC2B of (**C**) mice administered ɑ-IgG2b or ɑ-PDCA1 to deplete pDCs, or **(D)** WT or *Ifnar-/-* mice. (**G**) Bone marrow cells that were either a 1:1 mixture of CD45.1^+^ WT and CD45.2^+^ *Kdm5c^ΔItgax^* cells or CD45.2^+^ *Kdm5c^ΔItgax^* cells alone were transferred into lethally irradiated CD45.1^+^ WT mice. Proportions of splenic cDC subsets derived from CD45.1^+^ WT and CD45.2^+^ *Kdm5c^ΔItgax^*cells as a percentage of total cDCs were determined 7 weeks post-injection. Data are pooled from (**A**) two experiments (mean and s.e.m. of 10 to 11 mice per group), (**B**) two experiments (mean and s.e.m. of 7 to 8 mice per group), (**C**) two experiments (mean and s.e.m. of 10 mice per group), (**D**) three experiments (mean and s.e.m. of 14 mice per group), and (**E**) two experiments (mean and s.e.m. of 18 mice from mixed donor group and 9 mice from *Kdm5c^ΔItgax^*donor alone). Statistical significance was determined by (**A-D**) unpaired *t*-test or (**E**) one-way ANOVA. * p < 0.05, ** p < 0.01, *** p < 0.001, **** p < 0.0001

*Cre* expression under the control of the CD11c promoter is not exclusive to cDCs, and also includes expression in pDCs, as detailed above, along with some macrophages (36); therefore, to examine the effects of KDM5C deletion specifically in cDCs, we generated mice with KDM5C deleted in ZBTB46-expressing cells (*Kdm5c^ΔZbtb46^*). Comparison of cDC1 versus cDC2 populations in *Kdm5c^ΔZbtb46^* mice showed similar trends as *Kdm5c^ΔItgax^* mice (**Fig 3B**); however, the magnitude was less than that observed in the *Kdm5c^ΔItgax^*mice. Further, cDC2A and cDC2B proportions were not affected by *Kdm5c* deletion in *Kdm5c^ΔZbtb46^* mice (**Fig 3B**). We examined KDM5C deletion by western blot and found that KDM5C deletion was more efficient in *Kdm5c^ΔItgax^* DCs compared to *Kdm5c^ΔZbtb46^*DCs (**Supplementary Fig 3B**). There are several possibilities as to why these two models differ. *Cd11c-Cre* and *Zbtb46-Cre* are both expressed in the pre-cDC stage(36); however, the relative timing of their expression has not been accurately determined. In addition to differences in timing of Cre transgene expression and deletion efficiency, the difference in the cDC2 populations in the *Kdm5c^ΔZbtb46^*and *Kdm5c^ΔItgax^* mice could be a result of environmental differences due to KDM5C deletion in other cell types, since *Cd11c-Cre* expression is less restricted than *Zbtb46*-Cre.

Because KDM5C deficiency in *Cd11c*-expressing cells results in altered pDCs, we examined whether the altered pDCs contributed to the differences in cDC heterogeneity. We first depleted pDCs in wild type mice by administering 250 µg of anti-PDCA-1 antibody intraperitoneally every other day for one week *in vivo* (**Supplementary Fig 3C,D**). Depletion of pDCs resulted in a small but significant increase in the proportion of cDC2B relative to cDC2A (**Fig 3C**), but without the changes in cDC1 and cDC2A population sizes seen in the *Kdm5c^ΔZbtb46^* and *Kdm5c^ΔItgax^* mice (**Fig 3C; Supplementary Fig 3F)**. These results were reproduced in *Ifnar*-deficient mice, indicating that pDCs and type I IFN contribute moderately to changes in proportions of cDC2 subsets, but not cDC1s (**Fig 3D; Supplementary Fig 3G**). To further examine the effects of the environment on cDC heterogeneity, we generated bone marrow chimeric mice as in Fig 1B and 2B. We found that, similar to our findings for pDCs, the population dynamics of KDM5C-deficient cDCs were retained in the presence of WT hematopoietic cells (mixed chimera) and in the WT environment (**Fig 3E**), demonstrating that KDM5C deletion has both intrinsic and extrinsic effects on cDC population heterogeneity.

### KDM5C controls gene expression in cDC at homeostasis

We performed RNA-seq on control and KDM5C-deficient cDC1, cDC2A and cDC2B. KDM5C deletion resulted in the greatest effects on cDC1 (∼2000 differentially expressed genes) compared to ∼600 and ∼800 DE genes in cDC2A and cDC2B, respectively. KDM5C-deficient cDC1s displayed enrichment of transcripts encoding proteins in inflammatory pathways, cytokine pathways, defense to viral infection, immune cell activation and IRF regulated pathways (**Fig 4A,B, S4A, B, C**). These data are consistent with our previous work in BMDCs showing that KDM5C restrains DC activation (23). We performed transcription factor binding site analysis using Hypergeometric Optimization of Motif EnRichment (HOMER) on genes with significantly increased expression levels in KDM5C-deficient cDC1s compared to control, and found predominant enrichment of IRF family TF motifs (**Fig 4C**). This agreed well with our RNA-seq data which indicated the dysregulation of several *Irf* genes in the absence of KDM5C (**Fig 4D Heatmap**). Specifically, *Irf7*, *Irf1, Irf4* and *Irf2bpl* were significantly upregulated in KDM5C-deficient cDC1 compared to controls. IRF proteins are well known to participate in DC activation and activate inflammatory pathways, and therefore are likely key mediators of the increased gene expression observed in KDM5C-deficient cDC1.

**Figure 4 -.**
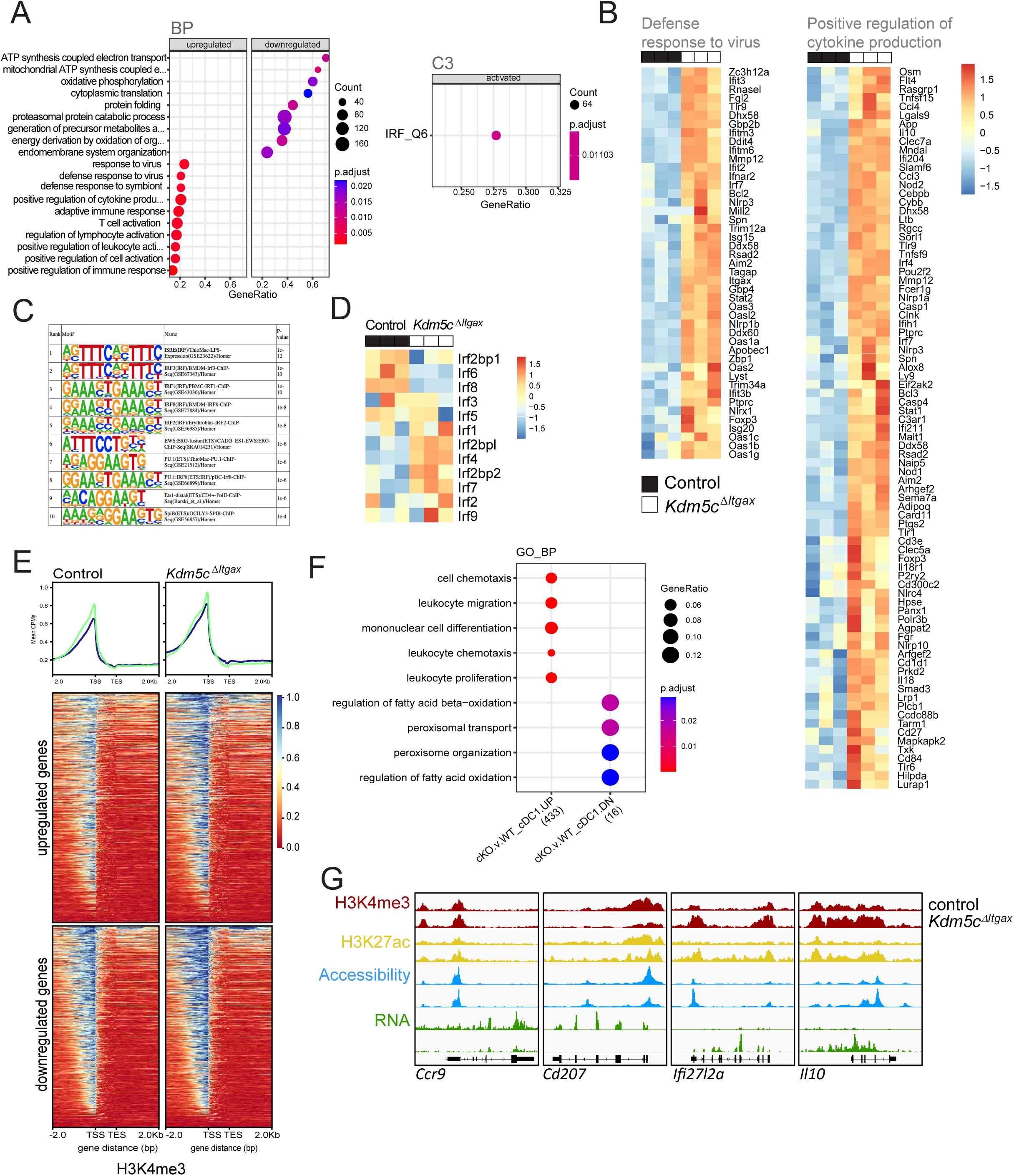
Disruption of epigenetic and transcriptional programming in cDC1 at homeostasis in the absence of KDM5C. RNA-seq analyses of control and KDM5C-deficient cDC1. Pathway analyses (**A**) and heatmaps (**B**) of differentially expressed genes. (**C**) HOMER analyses of promoters of genes upregulated in the absence of KDM5C. (**D**) Heatmap of Irf genes. (**E**) Heatmaps of H3K4me3 (CUT&RUN) segregated by direction of differential expression. (**F**) Pathway analysis of genes with differentially methylated regions. (**G**) IGV tracks showing H3K4me3, H3K27Ac, Accessibility (ATAC-seq) and RNA expression (RNA-Seq). Data is shown is combined from 3 biological replicates of each genotype.

KDM5C modifies chromatin structure by regulating the levels of H3K4me3, a histone PTM associated with active chromatin state and gene transcription. We examined H3K4me3 levels by Cleavage Under Targets and Release Using Nuclease (CUT&RUN)^19,26^ on ∼20K sorted cells of each subset. We found that global H3K4me3 levels increased in KDM5C-deficient cDC1s consistent with KDM5C being an H3K4 demethylase. The majority of the differentially methylated regions were found in intergenic regions (**Fig S4D**). To understand how changes in H3K4 trimethylation correspond to changes in gene expression in KDM5C-deficient DCs, we analyzed H3K4me3 levels in genes that were up or down regulated in our RNA-seq analysis (**Fig 4E**). As expected, we found increased H3K4me3 in genes that were upregulated in KDM5C-deficient cDC1s (**Fig 4E**). Surprisingly, we also found increased H3K4me3 associated with genes that showed decreased expression (**Fig 4E**). Regions with increased H3K4me3 were found in genes associated with immune cell migration, and proliferation (**Fig 4F**). Genes that were upregulated, such as *Ifi27* and *Il10*, were commonly associated with increased H3K4me3, chromatin accessibility (ATAC-seq) and H3K27ac (a marker of active chromatin) (**Fig 4G**). Genes such as *Cd207* that were downregulated had decreased chromatin accessibility and H3K4me3, whereas decreased expression of other genes such as *Ccr9* did not show corresponding changes in H3K4me3, chromatin accessibility or H3K27ac (**Fig 4G**). Together, these data support our model in which KDM5C restricts DC activation through the specific demethylation and inactivation of promoters of pro-inflammatory genes, including many IRF genes.

### KDM5C regulates OXPHOS gene expression and mitochondrial function in cDC1

As shown in **Fig 4E**, pathway enrichment analysis of our RNA-seq data identified the decreased abundance of transcripts associated with mitochondrial metabolism, including oxidative phosphorylation, in KDM5C-deficient cDC1s. Since bioenergetic metabolism is important for the function of DCs (37–40), we next examined the expression of genes encoding factors involved in oxidative phosphorylation and found that many genes were significantly decreased in KDM5C-deficient cDC1s (**Fig 5A**). To determine if altered expression of mitochondrial metabolism genes results in altered mitochondrial function, we first analyzed mitochondrial content and membrane potential using the mitochondrial dyes MitoSpy Green and TMRM, respectively. We found that with KDM5C deficiency, mitochondrial mass was lower, as was mitochondrial membrane potential (**Fig 5B)**. However, when graphed as a ratio, KDM5C-deficient cDC1 had increased membrane potential relative to mitochondrial mass. These differences were not due to changes in cell size (**Supplementary Fig 5A**).

**Figure 5 -.**
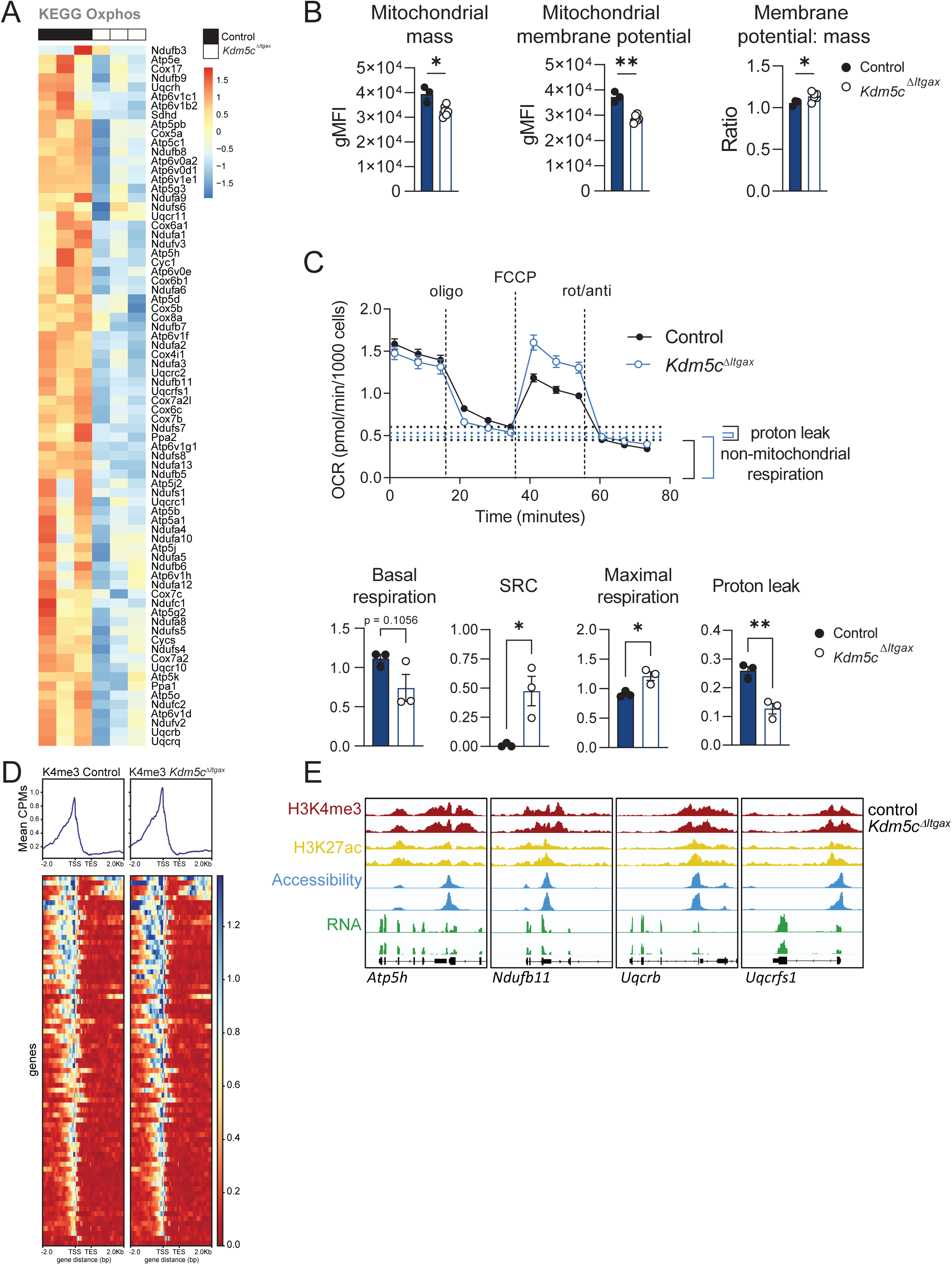
KDM5C regulates OXPHOS gene expression and mitochondrial function in cDC1. (**A**) Heat map of differentially expressed genes encoding factors from KEGG OXPHOS pathway by splenic cDC1 from control and *Kdm5c^ΔItgax^* mice. (**B**) Geometric MFI of MitoSpy Green (mitochondrial mass) and TMRM (mitochondrial membrane potential), and ratio of mitochondrial membrane potential to mass. (**C**) Oxygen consumption rate (OCR) measured over time of sorted control and *Kdm5c^ΔItgax^* cDC1 sequentially treated by oligomycin, FCCP, and rotenone/antimycin A (top). From this profile, several parameters were calculated (bottom): basal respiration (OCR of cells without drug treatment), spare respiratory capacity (SRC; maximal respiration minus basal respiration), maximal respiration (OCR following FCCP treatment), and proton leak (OCR following oligomycin treatment minus OCR following rotenone/antimycin A treatment). (**D**) Heatmap of H3K4me3 of genes shown in A. (**E**) IGV tracks showing H3K4me3, H3K27Ac, Accessibility (ATAC-seq) and RNA expression (RNA-Seq). Data is shown is combined from 3 biological replicates. Data are of one experiment representative of (**B**) two experiments (mean and s.e.m. of 3 WT and 6 *Kdm5c^ΔItgax^* mice), (**C**) two experiments (mean and s.e.m. of 3 replicates).

To assess whether these changes in mitochondria resulted in differences in cellular bioenergetics, we used a Seahorse bioanalyzer to measure cellular respiration. Surprisingly, we found that KDM5C-deficient cDC1s had a comparable baseline oxygen consumption rate (OCR) compared to the control, despite reduced mitochondrial mass (**Fig 5C**). Seahorse analysis uses the addition of several mitochondrial inhibitors to test the contribution of various mitochondrial processes to cellular OCR and provide insights into the causes of mitochondrial dysfunction. Oligomycin addition blocks the F_1_F_0_-ATPase and thus leftover OCR is due to proton leak into the mitochondrial matrix and/or non-mitochondrial respiration. KDM5C-deficient cDC1s displayed similar OCR following oligomycin treatment, however had increased non-mitochondrial respiration, suggesting that KDM5C-deficient cells have lower levels of proton leak (**Fig 5C**). Reduced proton leak can cause an increase in proton build up in the intermembrane space, and likely explains the enhanced mitochondrial membrane potential per mass that we observed using TMRM and MitoSpy Green. Since proton leak occurs passively or actively through uncoupling proteins (UCPs), we examined gene expression of UCPs in cDC1, and found a decrease in the expression of UCP2 (p.adj. 1.08E-06), potentially explaining the reduced levels of proton leak.

Spare respiratory capacity (SRC) is commonly assessed by measuring OCR following the addition of the mitochondrial uncoupler FCCP, which allows the release of protons from the inter membrane space, and results in maximal oxygen consumption as the mitochondria attempt to replenish membrane potential. We found there was a significant increase in SRC in the *Kdm5c^ΔItgax^*vs control cDC1s, which is again consistent with the increased TMRM/Mitospy Green ratio that we observed (**Fig. 5B,C**). Together, these data support a model in which, in the absence of KDM5C, changes in gene expression of OXPHOS genes and UCP2 result in lower overall mitochondrial mass, but increased mitochondrial membrane potential per mitochondria, and elevated SRC. These data also suggest that mitochondrial oxygen consumption is tightly coupled to ATP production in KDM5C-deficient cDC1s.

We examined H3K4me3 levels of OXPHOS genes with reduced expression in KDM5C-deficient cDC1. Surprisingly, we found that decreased expression was not associated with decreased H3K4me3 or decreased H3K27ac, a marker of active gene expression **Figs 5D, E**. We performed HOMER analyses on the same genes and found enrichment for YY1 motifs (**Fig S5B**). YY1 promotes the expression of mitochondrial respiration genes (41) through interaction with PGC-1ɑ. All together, these data strongly support a model in which KDM5C is a key regulator of mitochondrial function in cDC1s.

### KDM5C expression in DCs is required for T cell response to *Listeria* infection

In **Fig 4G** we showed that KDM5C results in altered gene expression of several IRF family members, including *Irf4* and *Irf8,* which encode lineage specifying transcription factors required for the generation and function of cDC1 and cDC2, respectively. We examined the expression of several lineage markers in cDC1s and found decreased expression of several cDC1-specific genes including *Irf8*, *Xcr1*, *Batf3*, *Cadm1*, and an increase in cDC2-specific genes including *Ltb*, *Cd4*, *Irf4*, *Itgam*, and *Tbx21*. Further, expression of *Tbx21*, which encodes cDC2A-specific transcription factor T-BET, was reduced in cDC2As to levels found in cDC2B (**Fig 6A**). Like the mitochondrial genes, changes in gene expression were not strongly associated with changes H3K4me3 or H3K27ac (**Fig 6B**). We however confirmed decreased IRF8 expression by flow cytometry (**Fig. 6C**). Thus, lineage specifying transcriptional programs are altered in the absence of KDM5C. Lineage-specific transcriptional programs are important for DC identity, differentiation, and function. To test whether the function of DCs is altered in the absence of KDM5C, we used the *Listeria monocytogenes* infection model in which proper cDC1 abundance and function are crucial for activating CD8^+^ T cells (42). Control and *Kdm5c^ΔItgax^* mice were infected with *Listeria monocytogenes* expressing ovalbumin (LM-OVA) for 7 days. Following infection with LM-OVA, the altered proportions of cDC and pDC subsets remained similar to those at homeostasis (**Fig 1A,B**), with *Kdm5c^ΔItgax^* mice exhibiting increased proportions of cDC1, cDC2B, and Ly6C^−^ pDCs, and decreased proportions of cDC2A and Ly6C^+^ pDCs in (**Supplementary Fig 6A,B**). Although there were no significant differences between control and *Kdm5c^ΔItgax^* mice in CD11a^+^CD49d^+^ antigen-experienced or CD44^+^ effector CD4^+^ T cells (**Fig 6F**), the proportion of *Listeria*-specific CD8^+^ T cells (tetramer^+^) was reduced in *Kdm5c^ΔItgax^* mice (**Fig 6D**). In addition, *Kdm5c^ΔItgax^* CD8^+^ T cells were less functional, as measured by decreased IFN-γ^+^ and IFN-γ^+^ TNF-ɑ^+^ polyfunctional CD8^+^ T cell populations (**Fig 6E**). Thus, KDM5C expression in DCs is necessary for CD8 T cell response to infection.

**Figure 6 -.**
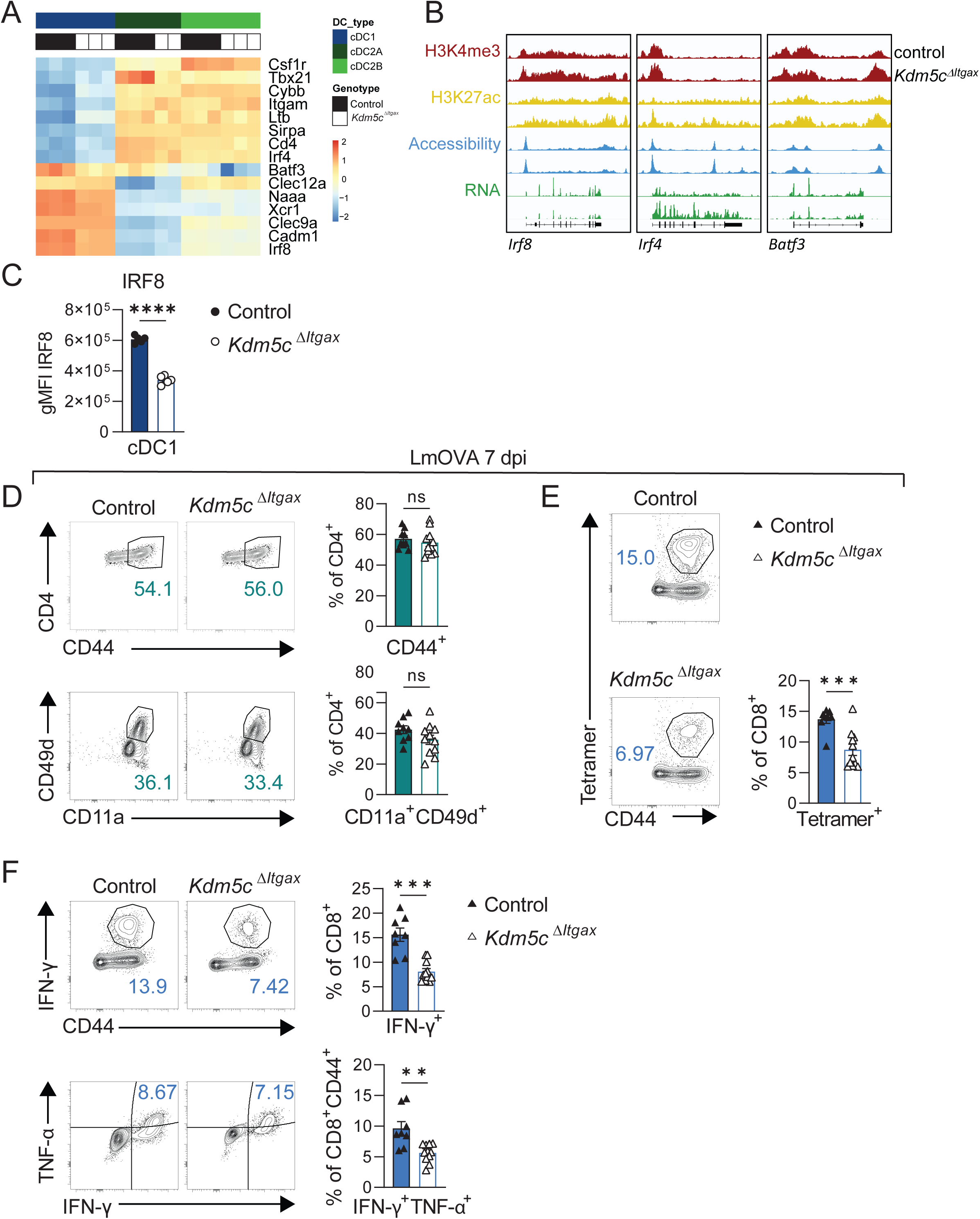
Alterations in DC lineage-specific gene expression and function in the absence of KDM5C. (**A**) Heat map of gene expression of markers specific to cDC subsets by cDC1, cDC2A, and cDC2B sorted from control and *Kdm5c^ΔItgax^* splenocytes. (**B**) IGV tracks showing H3K4me3, H3K27Ac, Accessibility (ATAC-seq) and RNA expression (RNA-Seq). Data is shown is combined from 3 biological replicates. (**C**) Geometric MFI of IRF8 in control and *Kdm5c^ΔItgax^* cDC1. Control and *Kdm5c^ΔItgax^* mice were infected with *Listeria monocytogenes* expressing ovalbumin (LmOVA) for 7 days, and assessed the frequencies of (**D**) CD44^+^ (top) and CD11a^+^CD49d^+^ (bottom) of CD4^+^ T cells (**E**) ovalbumin-specific (tetramer^+^) cells of CD8^+^ T cells, and (**F**) IFN-γ^+^ of CD8^+^ T cells (top) and IFN-γ^+^TNF-ɑ^+^ polyfunctional cells of CD8^+^CD44^+^ T cells (bottom). Data from (**C**) are of one experiment representative of three experiments (mean and s.e.m. of 5 mice per group), (**D-F**) are pooled from two experiments (mean and s.e.m. of 8 control and 11 *Kdm5c^ΔItgax^* mice). Statistical significance in (**C-F**) were determined by unpaired *t*-test.

## Discussion

Immune protection requires that immune responses be tailored to the infection or insult. DCs are among the first responders, and as a whole are specialized in antigen presentation and cytokine production. However, several subsets exist within the DC population whose functions are further specialized. Significant advances of the mechanisms that guide DC specification into these subsets are continuously being made (4,5,11,12,18,43). While several transcription factors have been demonstrated to be important for DC specification (17,18,44,45), our results show that histone modifying enzymes also influence DC cell fate. Our work implicates KDM5C as a key regulator of the balance of both pDC and cDC subtypes. In the absence of KDM5C, pDC and cDC population heterogeneity is altered and functions of both cell populations is impaired. KDM5C is not, however, required to generate a specific subset. Rather, it alters the proportions of subsets through modifying the epigenome and gene expression. Our data show that KDM5C is also important for functional responses of pDC and cDC, and that mice without KDM5C in DCs mount impaired CD8^+^ T cell response to *Listeria* infection.

Here, we identified a population of pDC that are Ly6C^−^, and are more prevalent in *Kdm5c^ΔItgax^* mice. Ly6C^−^ pDC are not well-studied, although one study also suggests that they produce less type I IFN than Ly6C^+^ cells (46). Recent studies have begun to highlight the heterogeneity and diverse functions of pDCs. Murine pDC-like (Lin^−^BST2^+^SiglecH^+^ZBTB46^+^) cells and transitional DCs (tDCs) (Lin^−^ CD11b^low^CD11c^+^SiglecH^+^CX3CR1^−^PDCA1^+^) resemble pDCs but share some characteristics and transcriptional similarities to cDCs(12,43). The pDC-like population has been described to serve as a progenitor pool for cDC2s(12). The specialized tDC exhibits an enhanced capacity for antigen presentation compared to pDCs and limited ability to produce type I IFNs, and can be further divided into CD11c^low^ tDC (Ly6C^+^), and CD11c high tDC (Ly6C^−^)(43). Although the Ly6C^−^ pDCs that were enriched in *Kdm5c^ΔItgax^* mice in our study had impaired type I IFN production, they were not exactly equivalent to the pDC-like or tDCs described in these other studies. Several lines of evidence from our work suggest that Ly6C^−^ pDCs are an immature pDC population: they have increased expression of cell cycle genes; their abundance is greater in the bone marrow, where pDC maturation occurs; and infection, IFN-β, and TLR-9 agonists induce a significant proportion of Ly6C^−^ pDC to become Ly6C^+^. Previous studies showed that CD4^+^ pDC are less migratory and produce lower levels of cytokines in response to stimulation compared to CD4^−^ pDC (32). However, in our data, Ly6C and CD4 expression did not co-segregate, suggesting they are not the same populations. This raises the possibility that there may be subsets within the Ly6C^−^ population. Further analyses at the single cell level are needed to understand the relationship between these subsets.

KDM5C-deficient Ly6C^+^ pDCs have an activated phenotype but are dysfunctional, which is a phenotype similar to that of exhausted pDCs (47,48). Exhausted pDCs have been found in models of persistent viral infections, have an activated phenotype, and fail to produce type I IFN upon CpG stimulation (48). KDM5C-deficient pDCs have an increase in expression of immune genes associated with activation, including *Ifi27*, *Ifi30*, MHCI and MHCII, as well as increased expression of several genes related to cytokine signaling. Our results suggest that KDM5C restrains the expression of immune response genes in pDCs, and lack of this restraint leads to dysfunctional responses to further stimulation.

We have shown that KDM5C promotes mitochondrial gene expression in bone marrow monocytes (BMM) (49), similar to what we observe here in cDC1s. However, KDM5C-deficient BMM have significant decreases in mitochondrial function, which is contrary to our observation that KDM5C-deficient cDC1s display enhanced mitochondrial coupling (reduced proton leak) and elevated spare respiratory capacity. The differences in the role of KDM5C in these cell types is not clear but could be related to the availability and use of metabolic fuels in experimental conditions. The KDM5A-C *Drosophila* ortholog KDM5 (Lid) promotes the transcription of genes important for mitochondrial function (50). However, KDM5 regulation of mitochondrial gene expression is mediated by the PHD3 domain, which is present in KDM5A/B but not KDM5C/D (50). How KDM5C then promotes transcription of these genes is not known. Positive regulation of gene expression by KDM5C has been linked to increased enhancer activity (by trimming H3K4me2/3 to H3K4me1) and co-activating gene expression (26,51). We found an enrichment of YY1 binding motifs, a transcription factor that promotes mitochondrial gene expression, suggesting that KDM5C function may be linked to this pathway. The implications of altered mitochondrial function for DC action are currently not clear; however, increased mitochondrial membrane potential to mass ratio has been associated with stress responses. Our results exemplify the importance of performing functional mitochondrial/metabolism assays to assess the consequences of changes in metabolic gene expression.

We found a strong IRF signature in genes dysregulated by cDC1 which likely is responsible for the increased inflammatory gene expression at steady state. IRF8 and IRF4 are key TFs involved in DC specification of cDC1 and cDC2, respectively. However, Kim *et al* show that IRF4 and IRF8 don’t specifically produce cDC2 and cDC1, but rather the amount of IRF protein determines their identity(16). While this study was focused on IRF4 and 8, it is not clear if other IRFs could also contribute to the tally of IRF that determine DC specificity. Deletion of IRF8 in committed cDC1s causes the cells to acquire a cDC2-like transcriptional signature and functional properties (52). We find that KDM5C fine-tunes lineage-specific gene expression beyond IRF4 and 8. KDM5C-deficient cDC1 have increased expression of several cDC2-specific genes and a concomitant decreased expression of cDC1-specific games.

The significance of DC subset heterogeneity and how changes in heterogeneity alter inflammation and immunity is unclear. Because different subsets have specialized function, it is likely that the composition of the DC population impacts the efficacy and efficiency of immune responses. The balance of DC subsets varies among mouse strains, individuals, lifespan and during inflammation(5,53–56). Our findings demonstrate that there are factors such as KDM5C that influence the balance of DC subtypes, along with the ability of the DC population as a whole to respond to infection. How KDM5C impacts DC specification and whether its expression or function is regulated during infection, aging, or across strains warrants further examination. Additional investigation into how the balance of subsets within the DC population affects immune responses will also provide further insight into how these cell types interact and work together to orchestrate a fully competent and efficient immune response in real world infection settings.

## Experimental procedures

### Mice

The following mouse strains were purchased from Jackson Laboratory and used for experiments and generating mouse lines used in this study; C57BL/6J (000664), *Ifnar^−/−^* (032045), *Zbtb*-Cre (028538), *Itgax*-Cre (008068), and Ly5.1 (002014). *Kdm5c*-fl/fl mice were a gift from the laboratory of Dr. Y. Shi and generated as described in (26,49,57). *Kdm5c*-fl/fl mice were crossed to *Itgax* and *Zbtb* Cre strains to produce the conditional knockout animals used in this work. Control mice were Kdm5cfl/fl-Cre negative. All mice were bred and maintained at Van Andel Research Institute Vivarium. All procedures involving mice were completed under approved IACUC protocols. All mice used were sex and age matched.

### Tissue processing

Spleens were injected with HBSS (with Ca^2+^ and Mg^2+^) containing 1 mg/mL of collagenase D and 10 μg/mL of DNase I (Roche) and incubated for 20 minutes at 37°C, followed by mashing and an additional 20 minute incubation at 37°C. The cell suspension was passed through a 70 μm cell strainer, then spun down at 300 × g for 5 minutes. The cell pellet was resuspended in 1 mL of RBC lysis buffer (155 mM NH_4_Cl, 12mM NaHCO_3_, and 0.125 mM EDTA) for 2 minutes, followed by addition of 4 mL of complete medium. The cells are spun down at 300 × g for 5 minutes and resuspended at the desired volume and buffer for downstream use.

### Mouse models

#### pDC depletion

Mice were injected intraperitoneally with 250 μg of α-PDCA-1 or IgG2b antibody (Bio Xcell) in 100 μL of PBS every other day. Five days following the first injection, pDC depletion was confirmed in blood collected by retro-orbital bleed and stained for flow cytometry analysis. Following confirmation of pDC depletion, mice were sacrificed the next day and spleens were removed and processed to analyze by flow cytometry.

#### Bone marrow chimera

Ly5.1 mice were irradiated with two doses of 450 rad four hours apart. The next day, bone marrow from age-matched Ly5.1 mice (WT) and *Kdm5c^ΔItgax^* mice were extracted from the femurs and tibia. Cells were counted and resuspended to 25 × 10^6^ non-RBCs per mL of PBS. 200 μL (5 × 10^6^) of bone marrow cells from WT, *Kdm5c^ΔItgax^*, or a 1:1 mix of WT and *Kdm5c^ΔItgax^* were intravenously injected per irradiated Ly5.1 mouse. Mice were kept on drinking water containing 0.17 mg/mL of enrofloxacin (pH 3.0) for 2 weeks. Spleen and bone marrow were collected and processed 7 weeks post-injection to examine by flow cytometry.

#### LCMV

Mice were infected with 2 × 10^5^ PFU of LCMV Armstrong, which was diluted in PBS from a frozen stock and delivered intraperitoneally in a BSL2 biosafety cabinet. Infected mice were housed in a separate quarantine room and monitored for the indicated length of time until tissues were collected and processed.

### Listeria

Attenuated *Listeria monocytogenes* expressing ovalbumin was grown in Tryptic Soy Broth containing streptomycin for 2-3 hours at 37°C and 250 rpm. Once an OD600 of 0.6 to 0.8 was reached, the bacteria were pelleted and diluted in PBS to a concentration of 2 × 10^7^ CFU/mL. Listeria was administered intravenously at 2 × 10^6^ CFU per mouse in a BSL2 biosafety cabinet. Mice were housed in a separate quarantine room and monitored for 7 days until tissues were harvested for immunophenotyping. For ex vivo analyses, splenocytes from mice infected for 7 days with LM-OVA were harvested and plated at 2×10^6^/well in a 96-well non-tissue culture-treated plate. Cells were washed and stimulated in the presence of OVA^257-264^ (1 µg/ml) (Anaspec), recombinant murine IL-2 200 U/ml (Peprotech) and 1x Brefeldin A (BioLegend) at a final volume of 200 µl for 5.5 hours at 37°C in a humidified incubator. Following stimulation, cells were washed and processed for flow cytometry.

### Flow cytometry

Samples were incubated with Fc block and eFluor 506 Fixable Viability Dye (ThermoFisher) in PBS, followed by an antibody cocktail prepared in wash buffer (PBS with 1% FBS, 1 mM EDTA, and 0.05% NaN_3_). For intracellular staining, cells were fixed with IC fixation buffer (eBioscience/ThermoFisher) for 30 minutes following surface staining, permeabilized using Permeabilization Buffer (eBioscience/ThermoFisher) and incubated for at least one hour with antibodies targeting intracellular proteins. Samples were acquired on the Cytek Aurora spectral cytometer and data analyzed using FlowJo v10. For mitochondrial staining, cells were plated and warmed to 37°C before staining. Mitochondrial dyes MitoSpy Green (BioLegend, 424806) and TMRM (ThermoFisher Scientific, T668) were prepared at 2x in HBSS, warmed, and added 1:1 to plated cells. Plates were incubated at 37°C for 20 minutes, washed, and surfaced stained. Cell sorting was performed on a Symphony S6 (BD Biosciences) or Moflo Astrios EQ Sorter (Beckman Coulter).

### IFN-ɑ enzyme-linked immunosorbent assay (ELISA)

Ly6C^+^ and Ly6C^−^ pDCs that are PDCA-1^+^B220^+^CD11b^−^CD11c^int^ were sorted from *Kdm5c^ΔItgax^* or control spleens and samples were pooled together for biological replicates of 15,000 cells per well. Cells were resuspended in 100 µL of complete medium (RPMI 1640 and 10% FBS, 2 mM L-glutamine, 100 U/mL penicillin/streptomycin, 0.01% β-mercaptoethanol) prior to plating, and 2X treatment was added to each well in 100 µL of complete medium. Cells were treated with CpG ODN 1585 to stimulate IFN responses (Invivogen; catalog # tlr-kit9m). After treatment for 18 hours, supernatant was collected and IFN-ɑ measured by ELISA according to manufacturer’s instructions (Invitrogen; catalog # BMS6027)

### In vitro DC differentiation with FLT3L

Mouse tibia and femur were washed with 70 % EtOH followed by PBS, and bone marrow was extracted. 2 × 10^5^ bone marrow cells (not counting RBC) were seeded per well in a 96-well plate in RPMI-1640 containing 10% FBS (NuSerum), 2 mM L-glutamine, 100 U/mL of penicillin/streptomycin, 0.55 μM β-mercaptoethanol, and 100 ng/mL FLT3L (Peprotech), with or without 50 U/mL IFN-β (PBL Interferon Source). After 3 days, 2.5 × 10^5^ mitomycin-treated OP9-DL1 cells (kindly gifted by Boris Reizis laboratory) were seeded in a tissue culture-treated 96-well plate (58). The OP9-DL1 cells were allowed to settle for approximately 2 hours prior to transferring over the bone marrow cells with replenished media with 100 ng/mL FLT3L with or without IFN-β as before. After 4 additional days, cells were stained to analyze by flow cytometry.

### Western blot

Protein lysates from FACS-sorted cDCs from control and *Kdm5c^ΔItgax^* or *Kdm5c^ΔZbtb46^* spleens were prepared using CHAPS lysis buffer (Cell Signaling, 9852S) with protease inhibitors (Roche, 11836170001), quantified with Pierce 660 nm Protein Assay Reagent (ThermoFisher Scientific; 1861426), and mixed with Laemmli sample buffer (BioWorld 10570021). 25 µg of protein per sample was loaded into 4-20% pre-cast polyacrylamide gels (Bio-Rad) and proteins were transferred to methanol-activated polyvinylidine fluoride membranes. Membranes were blocked for 1 hour using 5% milk in Tris-buffer saline (TBS) containing 0.1% Tween-20 (TBS-T), followed by overnight incubation at 4°C with anti-KDM5C antibody (1:1000; Abcam; Ab194288) in 5% milk in TBS-T, or 1 hour at room temperature with anti-β-actin antibody (1:1000; Cell Signaling; 4967S) in 5% bovine serum albumin (BSA) in TBS-T. Following washes with TBS-T, membranes were incubated with anti-rabbit horseradish peroxidase-linked antibody (Cell Signaling; 7074S) at a dilution of 1:4000 in 5% milk for 2 hours at room temperature for KDM5C, or 1:10 000 in 5% milk for 1 hour at room temperature for β-actin. Blots were developed by enhanced chemiluminescence using SuperSignal West Dura Extended Duration Substrate (ThermoFisher Scientific; 34075) and Bio-Rad ChemiDoc MP Imaging System.

### Metabolic assay

Several metabolic parameters were assessed in sorted cDC1 using the XF Cell Mito Stress Test and Seahorse XFe96 Analyzer (Agilent). cDC1s were labeled with PE-XCR1 antibody for positive selection using magnet after pan-DC enrichment (Miltenyi). The cell plate was coated with poly-D-lysine, then 200,000 cells were seeded per well in XF RPMI medium containing 10 mM glucose and 2 mM L-glutamine, with pH adjusted to 7.4. The plate was then incubated at 37°C in a non-CO2 incubator for 1 hour prior to the assay. Cells were sequentially treated with oligomycin (1.5 µM), FCCP (3 µM), and rotenone/antimycin A (0.5 µM), and oxygen consumption rate (OCR) was measured. At the end of each assay, cells were stained with Hoescht stain (20 µM; ThermoFisher Scientific) for 15 minutes at 37°C, then imaged using a Cytation imaging reader (Cytek) to count cells. OCR measurements were normalized by cell number.

### ATAC-seq

Libraries for ATAC-seq were prepared from nuclei of 25,000 sorted cells using the Omni-ATAC protocol in which mitochondrial DNA is depleted, enabling increased read-depth on genomic DNA(59,60). Transposition and amplification reactions were performed using the Nextera XT DNA Library preparation kit (Illumina cat# 15032354) along with IDT for Illumina adapters. Libraries were size-selected using Ampure XP beads (cat# A63881) and sequenced 50 bp, paired-end on an Illumina NovaSeq6000 sequencer. The data were trimmed using TrimGalore v0.6.0 (https://github.com/FelixKrueger/TrimGalore) to eliminate low-quality bases, and subsequently aligned to the mouse reference genome GRCm38 using bwa mem v0.7.17(61). Duplicates were marked using SAMBLASTER v0.1.24 (62), and low-quality bases filtered using SAMtools v1.9 (63). The bamCoverage tool in deepTools v3.4.3 (64) was utilized to generate BigWig files, excluding regions blacklisted in the ENCODE v2 blacklist(65). Furthermore, deepTools was employed to create heatmaps that visualize the distribution of chromatin accessibility levels surrounding gene regions. Differential accessibility analyses were performed using diffbind version 3.8.4 (66).

### RNA-seq

15,000 cells were FACS sorted into 1.5ml tubes containing 350ul lysis buffer. (Norgen Biotek buffer RL) RNA was extracted using Single Cell RNA Purification kit from Norgen Biotek (cat# 51800) and quantified using qubit HS RNA Assay kit (cat# Q32852). RNA libraries were generated using Takara SMARTer Stranded total RNA-seq Kit v3 (cat# 634487) and sequenced 50 bp, paired-end on an Illumina NovaSeq6000 sequencer. Reads were aligned to the mm10 genome and ERCC sequences using Takara’s CogentAP v1.5, specifying the ‘Strnd_UMI’ kit configuration. The deduplicated (via UMIs) counts were imported into R v4.1.0 for further analysis. Genes with >2 counts in at least 2 samples were retained. Differential expression for pairwise contrasts was tested using DESeq2 v1.32.0(67) with a significance cutoff of 0.1 FDR; a model design of ‘∼ Group’ was used, where Group represents unique combinations of genotype, treatment and cell type. Log fold changes were shrunken using the ‘lfcShrink’ function, with the parameter, “type=’ashr’” (68) and used to rank genes for GSEA analysis using clusterProfiler v4.0.5 (69)).

### CUT&RUN

Libraries for CUT&RUN-seq were prepared from 25,000 sorted cells(70). Transposition and amplification reactions were performed using the Nextera DNA Library preparation kit (Illumina) and sequenced 50 bp, paired-end on an Illumina NovaSeq6000 sequencer. Reads were trimmed using TrimGalore v0.6.0 with default parameters and aligned to the mm10 genome and decoy sequences, including viral sequences and cfMeDIP-seq spike-in sequences, (71,72)using bwa mem v0.7.17 (61). PCR duplicates were marked using SAMBLASTER v0.1.24 (62). Only high-confidence and properly-paired alignments were retained using samtools view with parameters, “-q 30 -f 2 -F 2828” (v1.9)(73). For peak calling, duplicate alignments were removed and processed with MACS2 v2.2.7.1(74) with the parameters, “-f BAMPE -g mm --keep-dup ‘all’”. Called peaks were filtered to remove ENCODE blacklist v2 regions(65). Bigwig files for visualization was produced using DeepTools v3.4.3(64); bamCoverage was run with the parameters, “--binSize 10 --extendReads --normalizeUsing ‘CPM’ -- samFlagExclude 1024 --samFlagInclude 64”. Bigwig files were combined across replicates by finding the mean using WiggleTools v1.2.11(75) and wigToBigWig from UCSC tools. Coverage heatmaps were generated using Deeptools v3.4.3.

Differential PTM was tested using DiffBind v3.2.7(76). For read counting, ‘dba.count’ was run with the parameters, ‘summits=200, bUseSummarizeOverlaps = TRUE’, with SummarizeOverlaps configured to paired-end mode. For sample normalization, ‘dba.normalize’ was run with the parameters, ‘normalize = DBA_NORM_NATIVE, background = TRUE’. The ‘dba.analyze’ step was run with the parameters, ‘bBlacklist=FALSE, bGreylist=FALSE’. Pairwise contrasts between different combinations of genotype and cell type were tested using an FDR cutoff of 0.1. Significant peaks were annotated to their nearest gene using ChIPSeeker v1.28.3(77), considering −3000 to +500 as the promoter region. For pathway enrichment analysis, peaks labeled as “Distal Intergenic” were removed and peaks were separated into up and down-regulated. Overlapping genes were tested using hypergeometric tests as implemented in clusterProfiler v4.0.5.

### Heatmaps

Heatmaps were generated using pheatmap version 1.0.12 package in R (version 4.4.2).

### HOMER Analysis

The findMotifs.pl script in HOMER(78) version 4.11 was utilized to identify transcription factor binding sites (TFBS) using gene lists obtained from transcriptomics differential expression analyses. The search criteria for TFBS included lengths ranging from 8 to 10 bases, and their locations were restricted to within 2000 bases upstream and 100 bases downstream of the transcription start site (TSS).

## Supplementary figure legends

cDC/pDC

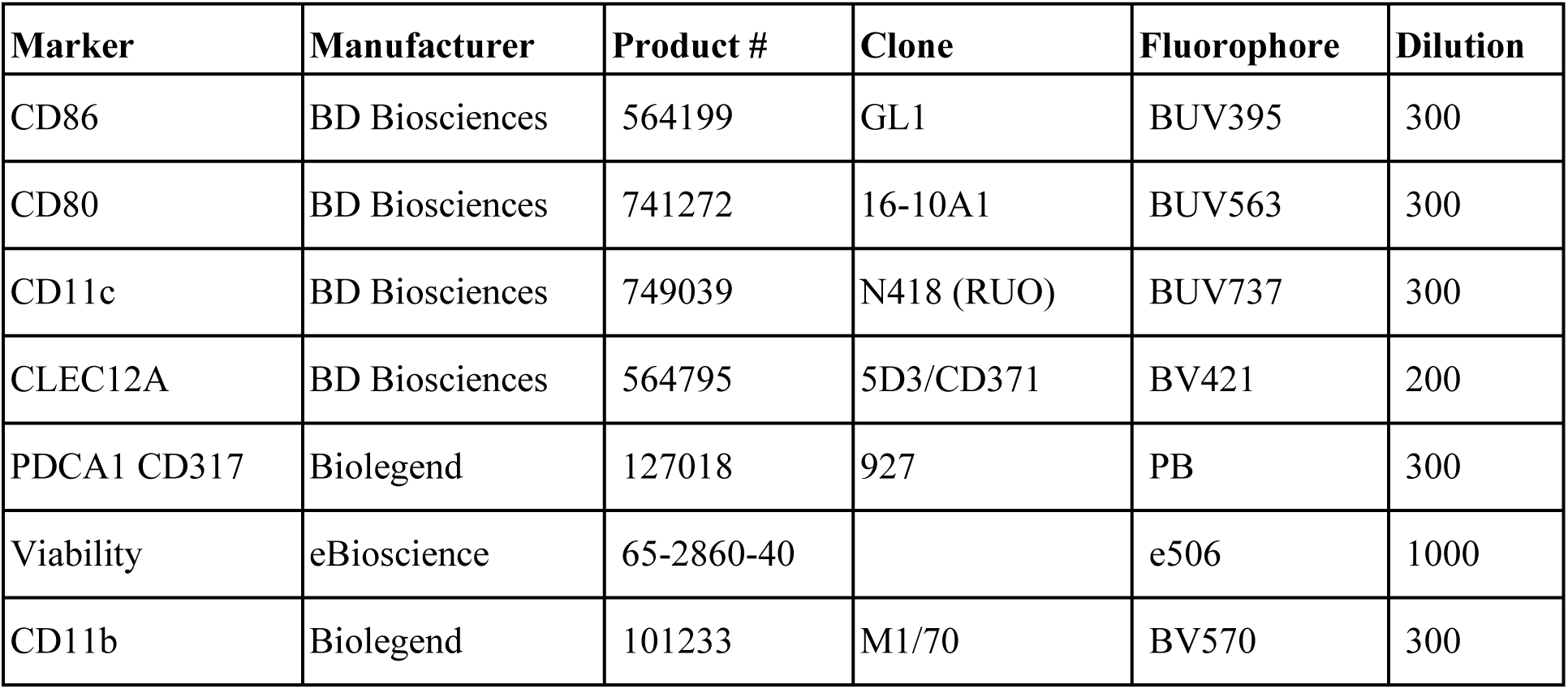
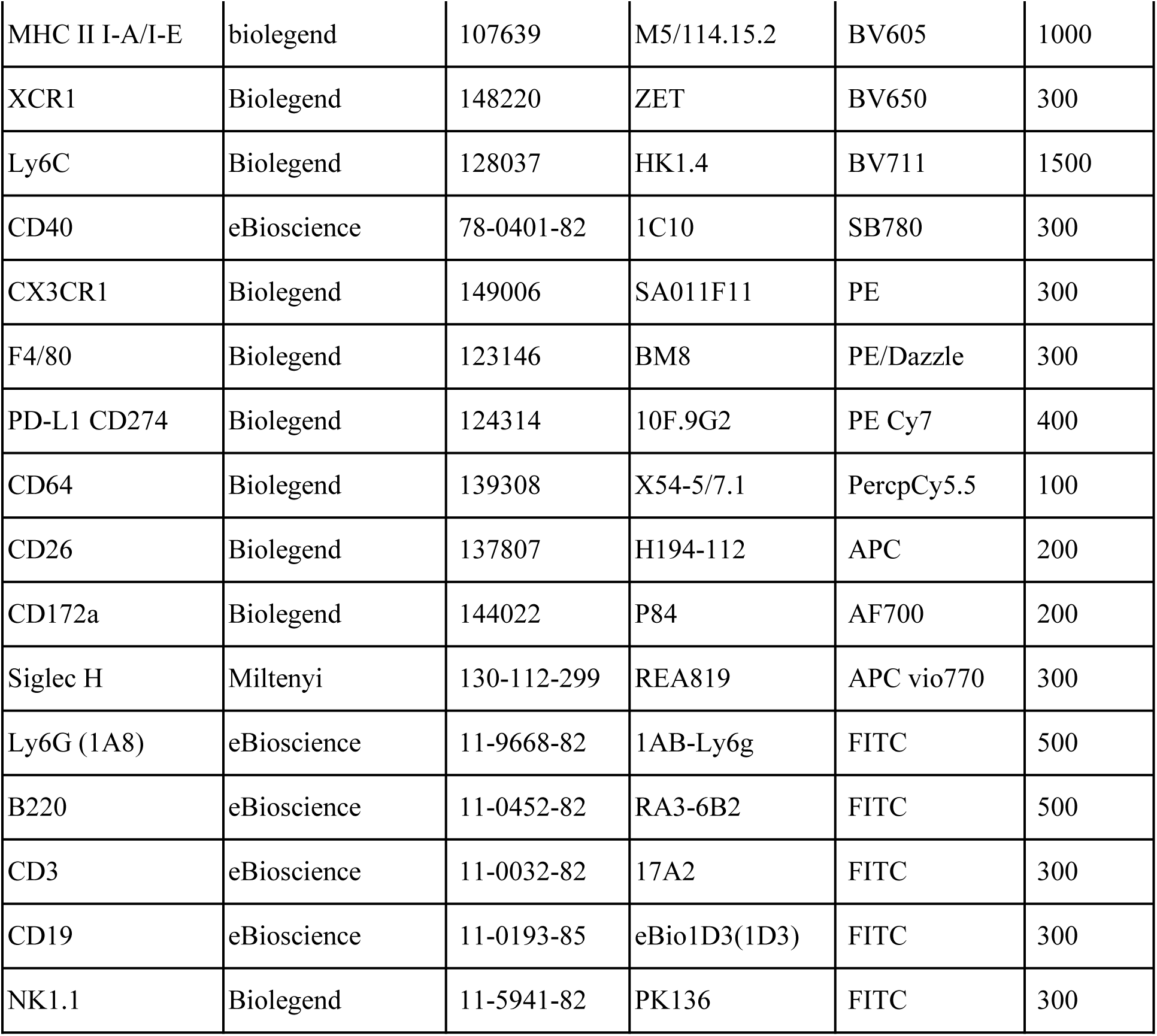

| Marker | Manufacturer | Product # | Clone | Fluorophore | Dilution |
| --- | --- | --- | --- | --- | --- |
| CD86 | BD Biosciences | 564199 | GL1 | BUV395 | 300 |
| CD80 | BD Biosciences | 741272 | 16-10A1 | BUV563 | 300 |
| CD11c | BD Biosciences | 749039 | N418 (RUO) | BUV737 | 300 |
| CLEC12A | BD Biosciences | 564795 | 5D3/CD371 | BV421 | 200 |
| PDCA1 CD317 | Biolegend | 127018 | 927 | PB | 300 |
| Viability | eBioscience | 65-2860-40 |  | e506 | 1000 |
| CD11b | Biolegend | 101233 | M1/70 | BV570 | 300 |
| MHC II I-A/I-E | biolegend | 107639 | M5/114.15.2 | BV605 | 1000 |
| XCR1 | Biolegend | 148220 | ZET | BV650 | 300 |
| Ly6C | Biolegend | 128037 | HK1.4 | BV711 | 1500 |
| CD40 | eBioscience | 78-0401-82 | 1C10 | SB780 | 300 |
| CX3CR1 | Biolegend | 149006 | SA011F11 | PE | 300 |
| F4/80 | Biolegend | 123146 | BM8 | PE/Dazzle | 300 |
| PD-L1 CD274 | Biolegend | 124314 | 10F.9G2 | PE Cy7 | 400 |
| CD64 | Biolegend | 139308 | X54-5/7.1 | PercpCy5.5 | 100 |
| CD26 | Biolegend | 137807 | H194-112 | APC | 200 |
| CD172a | Biolegend | 144022 | P84 | AF700 | 200 |
| Siglec H | Miltenyi | 130-112-299 | REA819 | APC vio770 | 300 |
| Ly6G (1A8) | eBioscience | 11-9668-82 | 1AB-Ly6g | FITC | 500 |
| B220 | eBioscience | 11-0452-82 | RA3-6B2 | FITC | 500 |
| CD3 | eBioscience | 11-0032-82 | 17A2 | FITC | 300 |
| CD19 | eBioscience | 11-0193-85 | eBio1D3(1D3) | FITC | 300 |
| NK1.1 | Biolegend | 11-5941-82 | PK136 | FITC | 300 |

Sorting panel

| Marker | Manufacturer | Product # | Clone | Fluorophore | Dilution |
| --- | --- | --- | --- | --- | --- |
| MHCII | Biolegend | 107639 | M5/114.15.2 | BV605 | 600 |
| CLEC12A | Biolegend | 143405 | 5D3/CLEC12A | APC | 200 |
| XCR1 | Biolegend | 148224 | ZET | APCCy7 | 200 |
| CD3 | eBioscience | 11-0032-82 | 17A2 | FITC | 200 |
| CD19 | eBioscience | 11-0193-85 | eBio1D3(1D3) | FITC | 200 |
| NK1.1 | Biolegend | 11-5941-82 | PK136 | FITC | 200 |
| Ly6G | eBioscience | 11-9668-82 | 1AB-Ly6g | FITC | 200 |
| CD64 | Biolegend | 139316 | X54-5/7.1 | FITC | 200 |
| F480 | Biolegend | 123146 | BM8 | PE-Dazzle594 | 200 |
| CD11b | eBioscience | 45-0112-80 | M1/70 | Percp-Cy5.5 | 200 |
| CD11c | eBioscience | 25-0114-82 | N418 | Pe-Cy7 | 300 |
| CD172a | Biolegend | 144022 | P84 | AF700 | 200 |
| PDAC1 | Biolegend | 127023 | 927 | BV421 | 200 |
| B220 | Biolegend | 103241 | RA3-6B2 | BV650 | 200 |
| Ly6C | eBioscience | 12-5932-82 | HK1.4 | PE | 200 |
| viability |  |  |  | DAPI | 500 |

Tetramer panel

| Marker | Manufacturer | Product # | Clone | Fluorophore | Dilution |
| --- | --- | --- | --- | --- | --- |
| CD62L | BD Biosciences | 740218 | MEL-14 | BUV395 | 300 |
| CD4 | BD Biosciences | 741050 | RM4-5 | BUV496 | 300 |
| KLRG1 | BD Biosciences | 741586 | 2F1 | BUV661 | 300 |
| CD8a | BD Biosciences | 612898 | 53-6.7 | BUV805 | 300 |
| CD223 | eBioscience | 48-2231-80 | eBioC9B7W(C9B7W) | e450 | 300 |
| PD-1 | Biolegend | 135219 | 29F.1A12 | BV605 | 300 |
| CD44 | Biolegend | 103049 | 1M7 | BV650 | 600 |
| CD127 | Biolegend | 135037 | A7R34 | BV785 | 100 |
| CD3e | eBioscience | 45-0031-82 | 145-2C11 | PerCP Cy5.5 | 200 |
| B220 | eBioscience | 11-0452-82 | RA3-6B2 | FITC | 300 |
| Ly6G | eBioscience | 11-9668-82 | 1AB-Ly6g | FITC | 300 |
| NK1.1 | Biolegend | 108705 | PK136 | FITC | 300 |
| CD19 | eBioscience | 11-0193-85 | eBio1D3(1D3) | FITC | 300 |
| OVATET | Baylor College of Medicine | 108708 | Tetramer | PE | 250 |
| CD49d | Biolegend | 103626 | R1-2 | PE Dazzle | 600 |
| CD11a | Biolegend | 101122 | M17/4 | PECY7 | 800 |
| CD25 | eBioscience | 17-0251-82 | PC61.5 | APC | 300 |
| Viability | eBioscience | 65-0865-14 |  | APC e780 | 1500 |

Restim panel

| Marker | Manufacturer | Product # | Clone | Fluorophore | Dilution |
| --- | --- | --- | --- | --- | --- |
| KLRG1 | eBioscience | 58-5893-80 | 2F1 | AF532 | 300 |
| CX3CR1 | Biolegend | 149036 | SA011F11 | AF700 | 100 |
| CD8A | eBioscience | 47-0081-82 | 53-6.7 | APC Fire 750 | 300 |
| CD44 | BD Biosciences | 741921 | IM7 | BUV 805 | 300 |
| CD62L | Biolegend | 740218 | MEL-14 | BUV 395 | 300 |
| CD69 | BD Biosciences | 741063 | H1.2F3 | BUV 496 | 100 |
| CD127 | Biolegend | 135037 | A7R34 | BV785 | 100 |
| *LY108* | Biolegend | 134608 | 330-AJ | Pac Blue | 100 |
| Viability | Thermo Fisher | 65-2860-40 |  | e506 | 500 |
| TIM3 | Biolegend | 119727 | RMT3-23 | BV711 | 100 |
| CD279 | BD Biosciences | 749306 | RMPI-30 | BUV 737 | 100 |
| CD4 | eBioscience | 15-0042-83 | RM4-5 | PE-CY5 | 300 |
| IFNγ | eBioscience | 17-7311-82 | XMG1.2 | FITC | 100 |
| TNFα | eBioscience | 25-7321-82 | MP6-X1222 | PE-Cy7 | 200 |
| Tox | eBioscience | 46-4875-82 | Dan11mag | PE | 100 |
| TCF1 | eBioscience | 6709S | C63D9 | AF647 | 100 |
| GZMB | Biolegend | 372216 | QA16A02 | PE-Dazzle | 100 |
| TBET | Biolegend | 644817 | 4B10 | BV605 | 100 |

