## Supplementary figures and images for "The histone lysine demethylase KDM5C fine-tunes gene expression to regulate dendritic cell heterogeneity and function"

### S1

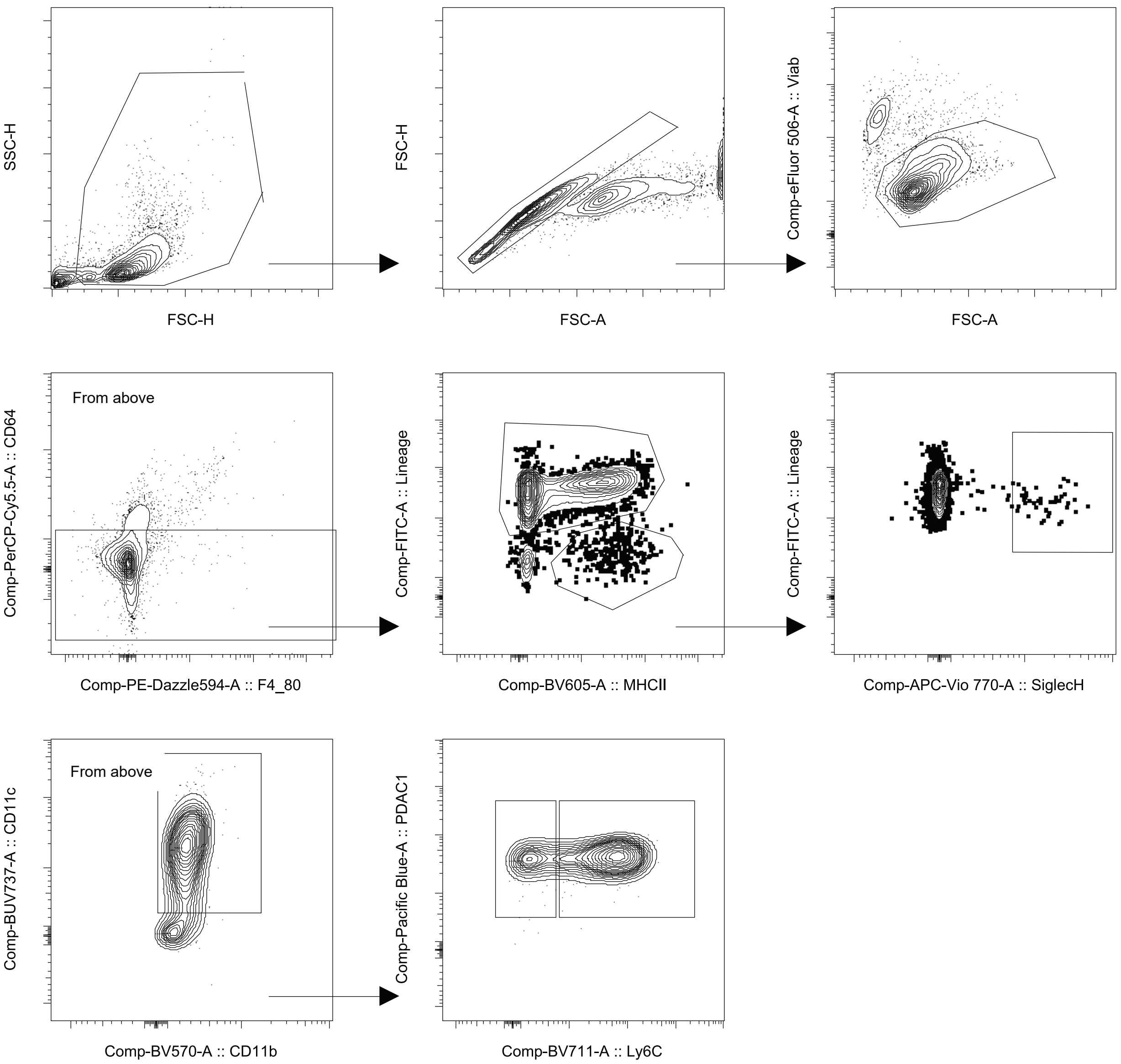


Supplementary Figure 1 - Flow Cytometry Gating Strategy for Ly6C+ and Ly6C- pDCs.
