## Supplementary material for "The histone lysine demethylase KDM5C fine-tunes gene expression to regulate dendritic cell heterogeneity and function": S2

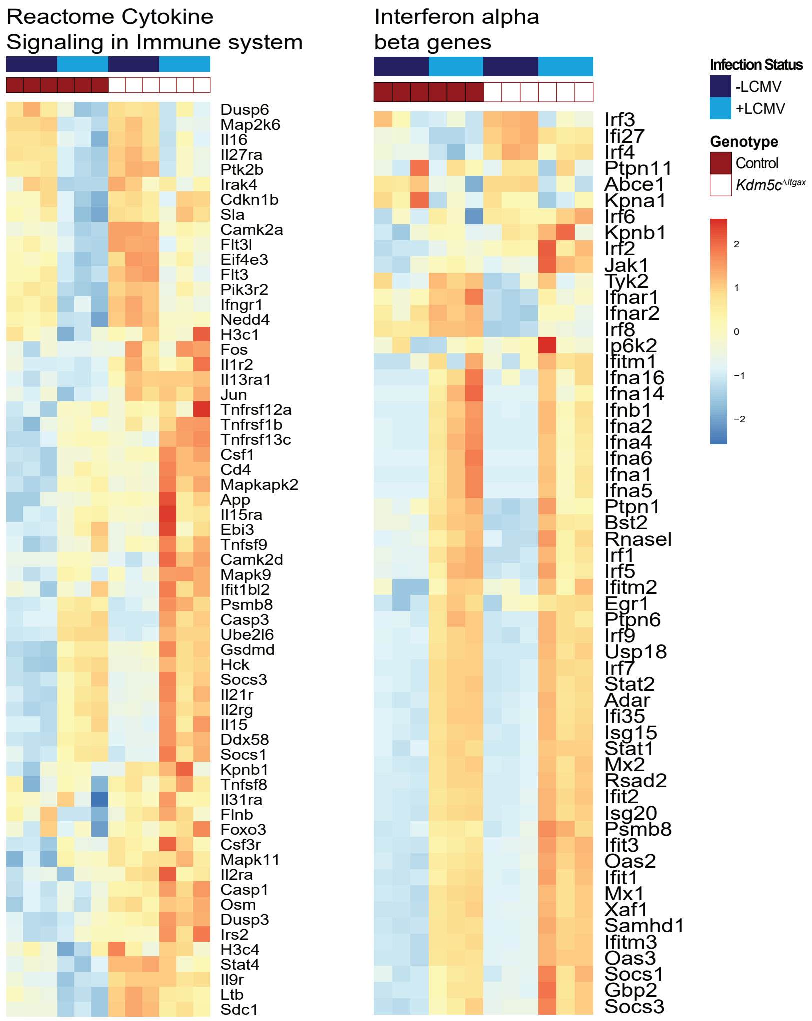


Supplementary Figure 2 - Related to Fig 2. Heat maps of differentially expressed genes in sorted control and KDM5C-deficient Ly6C+ pDCs at homeostasis and in response to LCMV infection (20 hrs).
