## Supplementary material for "The histone lysine demethylase KDM5C fine-tunes gene expression to regulate dendritic cell heterogeneity and function": S3

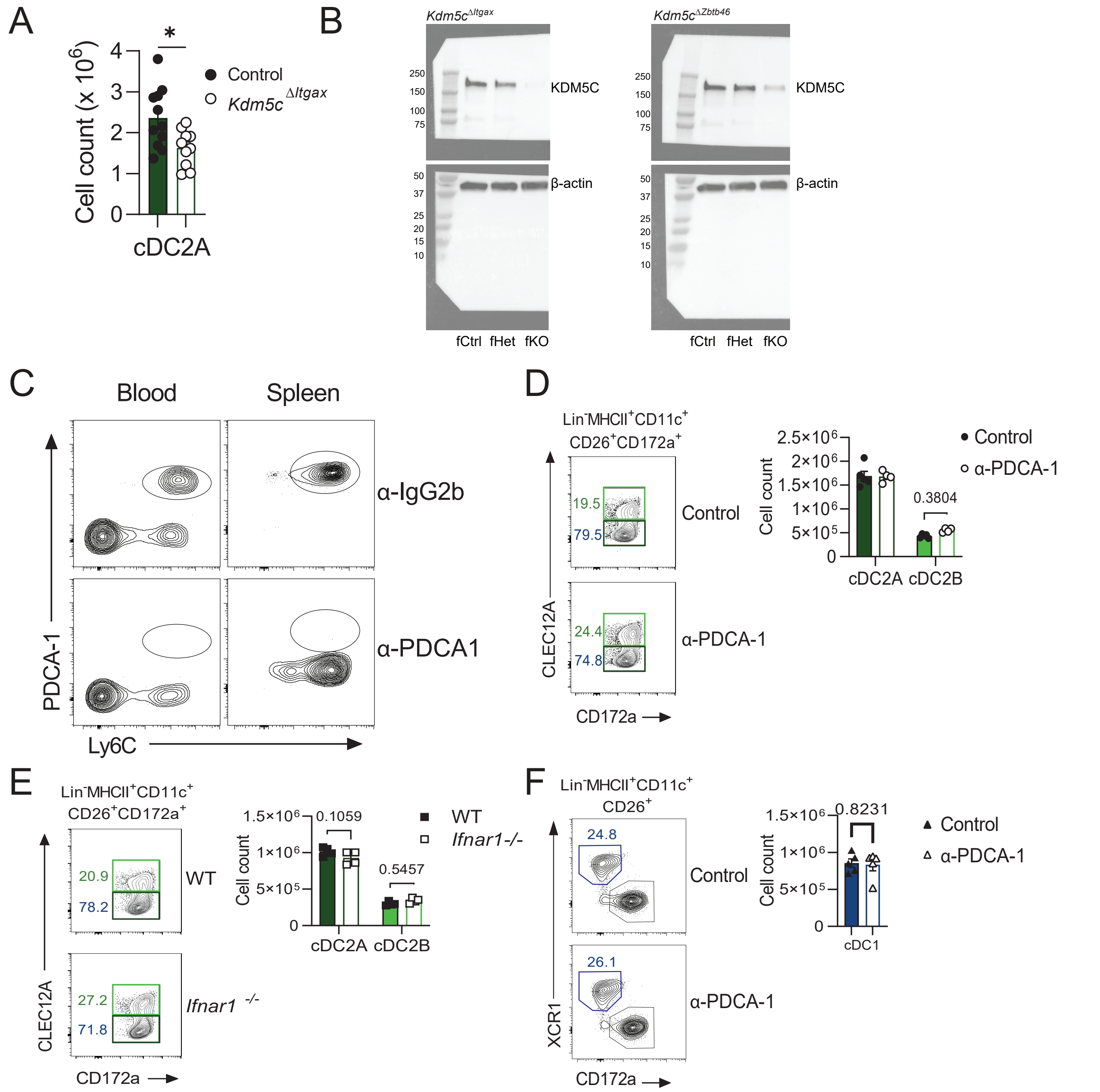


Supplementary Figure 3 - related to Fig. 3. (A) Cell count of splenic cDC2A from control and Kdm5cΔItgax mice. (B) Western blots of KDM5C and β-actin from sorted splenic cDCs from control, heterozygous, and conditional knockout mice with conditional deletion of Itgax (left) or Zbtb46 (right). Mice were administered α-IgG2b or α-PDCA1 as in Figure 3C and (C) pDC depletion in blood and spleen by α-PDCA1 was confirmed; (D) representative FACS plots , frequency of cDC2A and cDC2B as percentage of cDC2 , and cell count of cDC2A and cDC2B. (E) Representative FACS plots, frequency of cDC2A and cDC2B as percentage of cDC2, and cell count of cDC2A and cDC2B of WT and Ifnar-/- mice. (F) representative FACS plots, cell count of cDC1. Data in (A) two experiments (mean and s.e.m. of 10 to 11 mice per group), (B) one experiment representative of 2 experiments, (C) are one experiment representative of two experiments, (D) are of one experiment representative of two experiments (mean and s.e.m. of 4 to 5 mice per group), (E) are of one experiment representative of three experiments (mean and s.e.m. of 4 mice per group). Statistical significance was determined by unpaired t-test. * $p < 0.05$, ** $p < 0.01$
