## Supplementary material for "The histone lysine demethylase KDM5C fine-tunes gene expression to regulate dendritic cell heterogeneity and function": S4

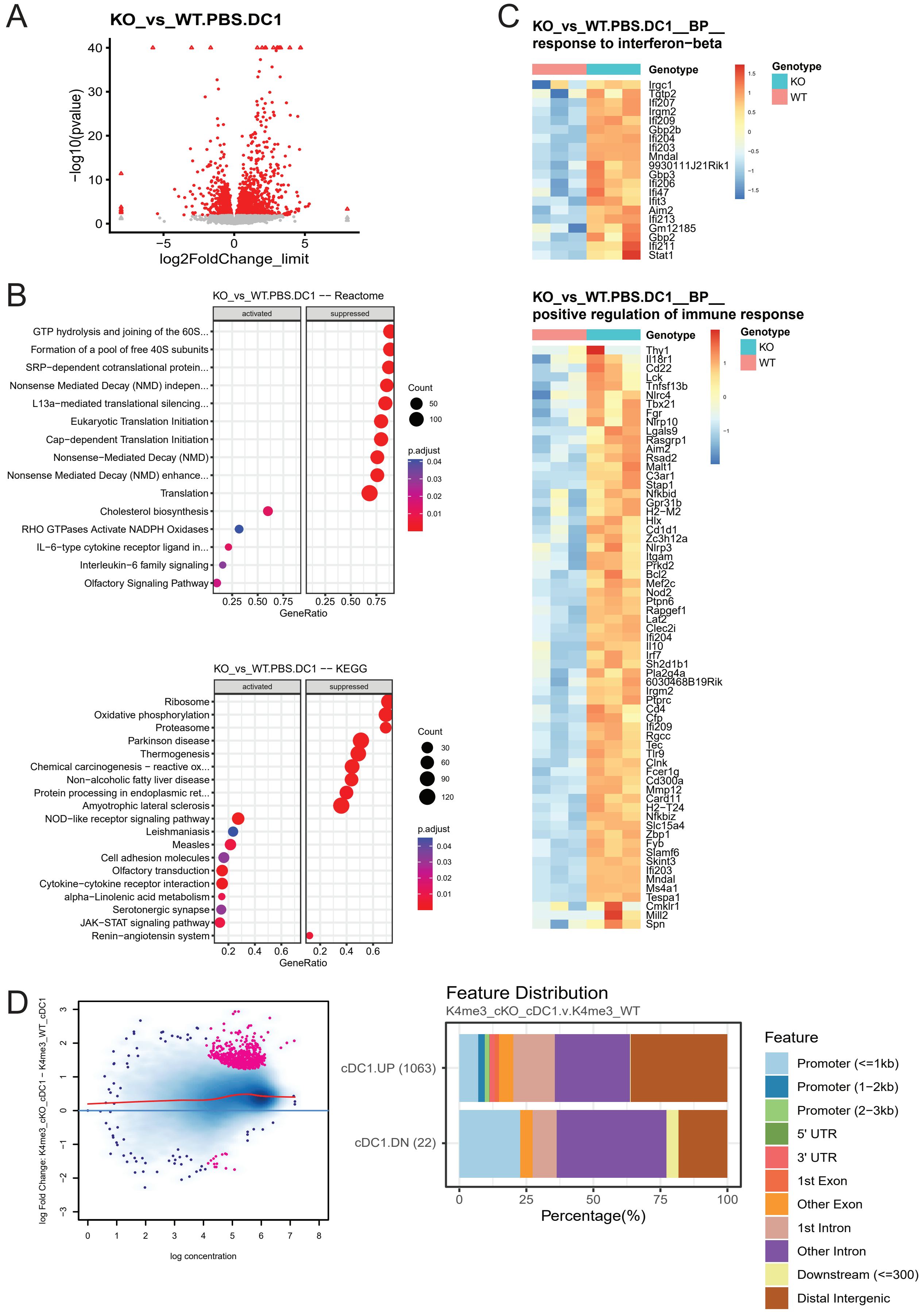


Supplementary Figure 4. related to Fig. 4. (A) Volcano plot of RNA-seq of the comparison of cDC1 cells from control and Kdm5cΔItgax. Red indicates significance. (B) Pathway analysis of differentially expressed genes from A. (C) Heat maps of differentially expressed genes encoding factors from GSEA BP of interferon response and immune activation by splenic cDC1 from control and Kdm5cΔItgax mice. (D) MA plot comparing H3K4me3 between control and KDM5C-deficient cDC1. Significantly different regions shown in pink. Feature distribution of the differentially methylated regions.
