## Supplementary material for "The histone lysine demethylase KDM5C fine-tunes gene expression to regulate dendritic cell heterogeneity and function": S5

A

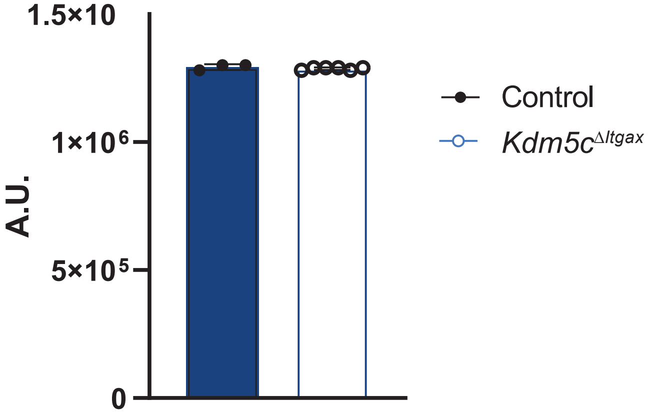


B

| Rank | Motif | Name | P-value |
|---|---|---|---|
| 1 | 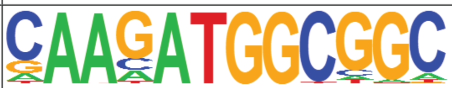 | YY1(Zf)/Promoter/Homer | 1e-5 |
| 2 | 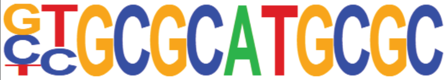 | NRF(NRF)/Promoter/Homer | 1e-5 |
| 3 | 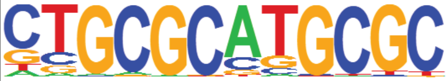 | NRF1(NRF)/MCF7-NRF1-ChIP-Seq(Unpublished)/Homer | 1e-4 |

Supplementary Figure 5 related to Fig. 5. (A) Cell size of cDC1 does not change with KDM5C loss. Forward scatter (FSC) of cDC1 in control and Kdm5cΔItgax mice as a measure of cell size. (B) HOMER analysis of OXPHOS genes shown in Fig 5A in KDM5C-deficient cDC1s. Data are of one experiment representative of (A) two experiments (mean and s.e.m. of 3 WT and 6 Kdm5cΔItgax mice).
