## Supplementary material for "The histone lysine demethylase KDM5C fine-tunes gene expression to regulate dendritic cell heterogeneity and function": S6

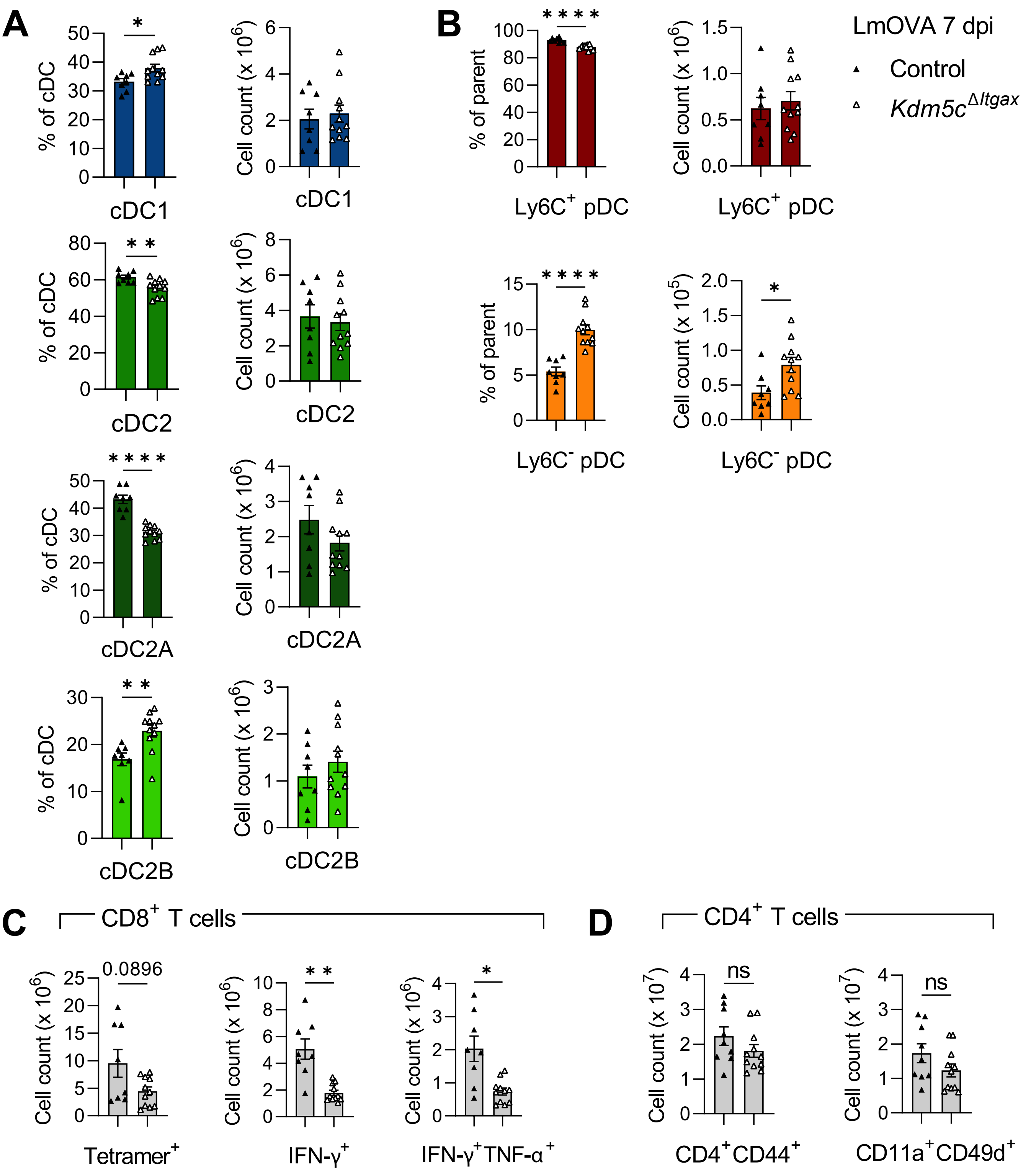


Supplementary Figure 6 - related to Fig. 6. Control and Kdm5cΔItgax mice were infected with LmOVA and examined 7 dpi. (A) Frequency of splenic cDC1, cDC2, cDC2A, cDC2B as a percentage of total cDCs (left column) and corresponding cell counts (right column). (B) Frequency of Ly6C+ and Ly6C- pDCs as a percentage of the parent population (Lin(B220)+SiglecH+CD11cintCD11b-) (left column) and corresponding cell counts (right column). Cell counts of (C) CD8+ T cells that are tetramer+, IFN-γ+, or IFN-γ+TNF-α+, (D) CD4+ T cells that are CD44+ or CD11a+CD49d+. Data are pooled from two experiments (mean and s.e.m. of 8 control and 11 Kdm5cΔItgax mice). Statistical significance was determined by unpaired t-test. * $p < 0.05$, ** $p < 0.01$, **** $p < 0.0001$
